# Alternative splicing changes are associated with pre-birth adaptation during lung development

**DOI:** 10.1101/2022.01.19.476886

**Authors:** Marta F. Fidalgo, Catarina G. Fonseca, Paulo Caldas, Alexandre A. S. F. Raposo, Tania Balboni, Ana R. Grosso, Francisca F. Vasconcelos, Cláudio A. Franco

**Affiliations:** Instituto de Medicina Molecular João Lobo Antunes, Faculdade de Medicina, Universidade de Lisboa, 1649-028 Lisboa, Portugal; UCIBIO – Applied Molecular Biosciences Unit, Department of Life Sciences, NOVA School of Science and Technology, NOVA University Lisbon, 2819-516 Caparica, Portugal; Department of Experimental, Diagnostic and Specialty Medicine, University of Bologna, Bologna, Italy; Instituto de Histologia e Biologia do Desenvolvimento, Faculdade de Medicina, Universidade de Lisboa, 1649-028 Lisboa, Portugal

**Keywords:** Lung development, alternative splicing, vascular endothelial growth factor A, endothelial cells, alveolar epithelial cells

## Abstract

Gas exchanges are ensured by lung alveoli, which are mainly composed by epithelial alveolar type 1 (AT1), alveolar type 2 (AT2) and capillary endothelial cells (ECs). Alveologenesis starts during late embryonic development and continues after birth and relies on extensive biochemical crosstalk between these cell types. How this crosstalk is modulated to anticipate and accommodate the radical changes occurring at birth is still unclear.

We investigated the alternative splicing (AS) changes occurring during lung development at the embryonic to postnatal transition by performing RNAseq of mouse lungs at distinct developmental stages. We found that most of the AS changes occur at the embryonic to postnatal transition. In addition, we identified hnRNP A1, Cpeb4 and Elavl2/HuB as putative splicing regulators of this transition. We show that the AS of a major pro-angiogenic chemokine, vascular endothelial growth factor A (VEGFA), is differentially regulated at this transition. Remarkably, we found that there is a switch from the predominance of *Vegfa 164* to *Vegfa 188* just before birth specifically in AT1 cells, whilst in other cell populations *Vegfa* does not undergo AS changes. Moreover, we identified a novel *Vegfa* isoform generated by the retention of intron 5, *Vegfa i5*.

Our results reveal a cell type-specific regulation of *Vegfa* AS that may constitute a pre-birth adaptation mechanism of the epithelial-endothelial crosstalk, which may be fundamental for the adaptation to breathing and may have implications for pathological conditions.

## Introduction

Respiration, the process of gas exchanges between the body and the environment, takes place at alveoli in the lung. The efficiency of gas exchanges is ensured by the functional specialization of the numerous cell types that compose the alveolus, including: epithelial alveolar type 1 (AT1) cells, thin and elongated cells that line the surface of each alveolus; epithelial alveolar type 2 (AT2) cells, which are sparsely distributed at the alveolar surface and secrete surfactant essential for alveolar inflation and deflation; and endothelial cells (ECs), which compose the interior surface of capillaries that tightly enwrap the alveolar epithelial layer, and promote gas exchanges with blood^1, 2^.

Alveologenesis starts during late embryonic development at E16.5 in mouse and continues for 3-8 weeks after birth. New alveoli form through branching morphogenesis and angiogenesis by coordinated growth and specialization of epithelial and endothelial cells, respectively ^2, 3^. The crosstalk between these cell types is mediated by multiple signalling pathways, such as VEGFA, WNT, FGF and HIPPO ^3–6^. For instance, secretion of VEGFA by epithelial cells during lung development regulates the expansion of the vascular network by binding to VEGF receptor 2 (VEGFR2) at the surface of ECs and triggering angiogenesis. Genetic deletion or pharmacological inhibition of VEGFA compromises lung alveolar epithelial development and capillary growth, leading to bronchopulmonary dysplasia, characterized by simplified alveoli and dysmorphic vasculature ^7–9^. Despite extensive research on lung development, how the communication between these cell types is modulated at the critical transition between embryonic to postnatal development is not fully understood.

Transcriptional and alternative splicing (AS) changes have previously been implicated in the regulation of multiple developmental processes ^10–13^. Previously published RNAseq and single-cell RNAseq studies from lungs at different developmental stages have enabled the comprehensive characterization of cell populations and the detailed study of gene expression changes occurring during lung development ^8, 14, 15^. Yet, due to the lack of sequencing depth, none of these approaches has allowed the study of AS, and thus, the knowledge regarding AS during lung development is limited. One of the few genes shown to undergo AS changes during lung development was *Vegfa* ^16–18^. *Vegfa* is composed of 8 exons and the most common *Vegfa* isoforms differ on the inclusion or exclusion of exons 6 and 7: *Vegfa 188* contains all eight *Vegfa* exons, *Vegfa 164* does not contain exon 6 and *Vegfa 120* does not contain exons 6 and 7. *Vegfa* isoforms are functionally distinct in terms of binding to the extracellular matrix, and in their potential to induce angiogenesis, EC proliferation, survival and vascular permeability ^19–22^. In addition, it has been shown that the relative proportion between *Vegfa 120*, *164* and *188* isoforms changes during lung development ^16–18^ and that loss of specific isoforms has a functional impact. Mice expressing exclusively *Vegfa 120* showed impaired lung development, whilst mice expressing exclusively *Vegfa 164* or *Vegfa 188* isoforms have no gross morphological defects ^23, 24^. Despite the relevance of *Vegfa* AS during lung development, the temporal dynamics and cell-specific expression of *Vegfa* isoforms along lung development remain poorly defined.

Here, we performed an unbiased genome-wide analysis of AS at embryonic and early postnatal stages of lung development and we identified that most of the AS changes occur at the transition from the pre- to postnatal life, suggesting that AS regulation may be associated with lung adaptation to birth. We validated our genome-wide analysis by focusing on *Vegfa* isoforms and we identified a cell type-specific switch in AS of *Vegfa* during lung alveologenesis.

## Results

### Genome-wide analysis reveals that AS changes occur at the transition from the pre- to post- natal period

To analyze the genome-wide AS changes occurring during lung development, we extracted and sequenced mRNA of whole mouse lungs at two embryonic (E15.5 and E18.5) and two postnatal (P5 and P8) stages **(Figure 1A)**. We performed 101 nucleotides (nt) paired-end RNAseq with triplicate samples and obtained an average of 59.8 million reads per sample **(Figure S1A).**

**Figure 1.**
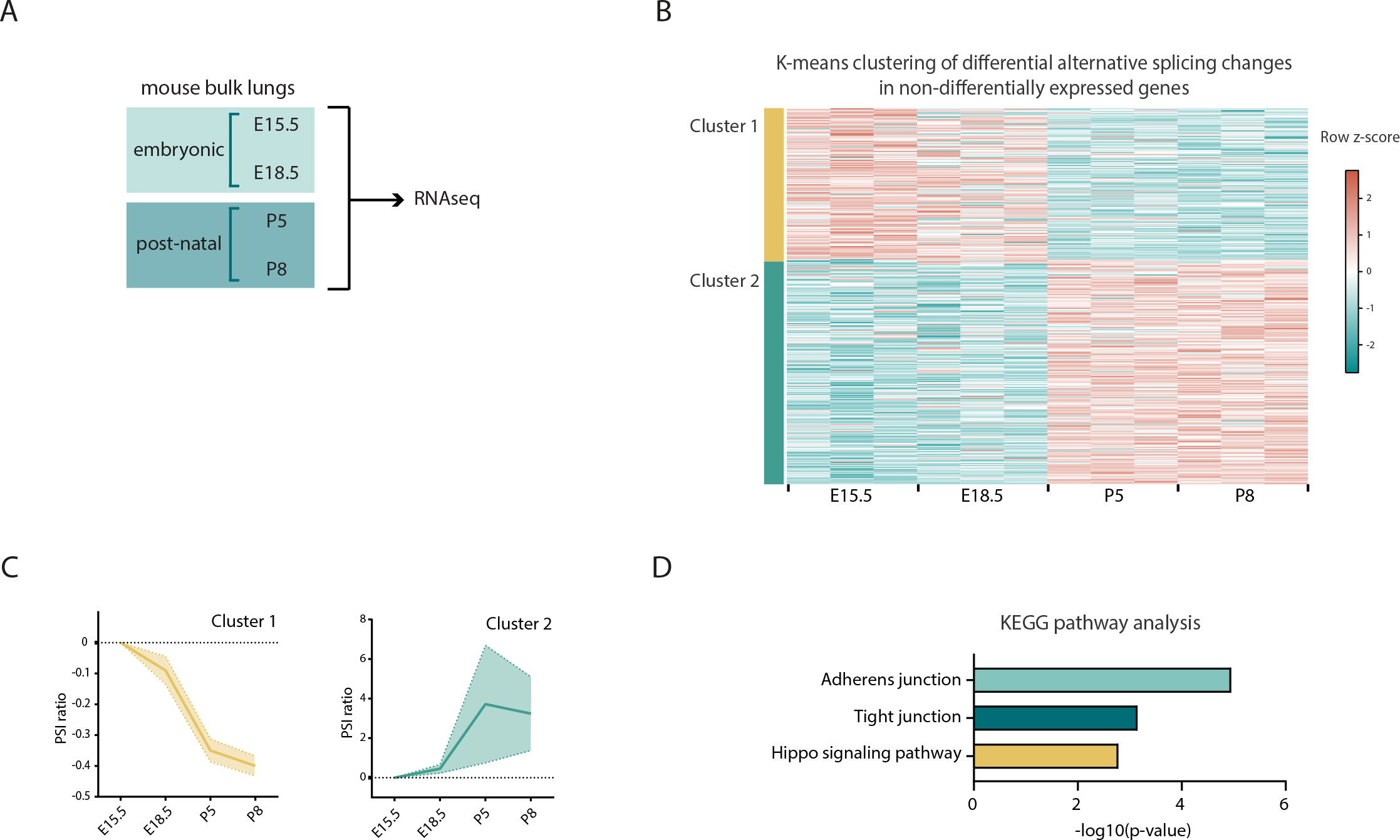
Genome-wide analysis of AS during lung development reveals that most splicing changes occur at the perinatal period. A. Schematic representation of workflow: Mouse lungs at different developmental time- points were collected. RNA was extracted and gene expression and splicing analysis were performed by RNAseq. B. Heatmap representing the K-means clustering of differentially AS events in at least one pair-wise comparison between time-points associated with genes that do not undergo differential gene expression in the same comparison (K=3). Cyan and coral represent decreased and increased PSI (percent spliced in), respectively, relative to the mean of each AS event across the time course (row z-score calculated from logit(PSI)). **Table S5** contains all AS events associated with each cluster. C. AS dynamics of the three kinetic clusters represented by mean ratio (PSI) (bold line) and 95% confidence interval (shaded area), data was centered in zero. D. Enrichment of KEGG terms associated with DAS events associated with non-differentially expressed genes. **Table S6** contains the genes associated with term and associated statistics.

To assess the quality of our datasets, we analysed the expression of a set of known lineage-specific markers **(Figure S1B)**. In accordance with what has been shown before ^25, 26^, our data shows that the epithelial progenitor marker Sox9 progressively decreases, while markers for AT1 (Aqp5, Pdpn) and AT2 (Sfptc, Sfptd) epithelial cells increase over the same developmental period. Also, the levels of EC-specific markers - Sox17, Pecam1 and Cdh5 - increase during development, as the lung progressively becomes more enriched in blood vessels. In addition, Car4 expression increases from E18.5 to P5, concordant with previous findings showing that Car4-positive ECs, a specialized type of alveolar capillary ECs (aCap), is specified just before birth at E19.5 ^8^. It has also been shown that, after birth, the immune cells repertoire that populates the lungs undergoes profound changes: Embryonic macrophages (Ncapd2-positive) are eliminated and postnatal lungs become enriched in T lymphocytes (Cd3e-positive), neutrophils (Retnig-positive) and several subtypes of macrophages (Itgax-, C1qa-, Plac8- positive) ^27^. Concordantly, in our datasets, the expression of Ncapd2 decreases at birth, while the expression of all these other immune cell markers increases at birth. Thus, this analysis supports that the obtained RNAseq datasets are of good quality.

To identify AS events that change during lung development, we used the Vertebrate AS database (VastDB) and Vertebrate AS Transcription Tools (VAST-TOOLS) ^28, 29^. We evaluated differential alternative splicing (DAS) between each pair of the above-mentioned perinatal time-points. We obtained 460 DAS events associated with 355 genes **(Table S1)** that showed a statistically significant variation in at least one of the pairwise comparisons (|ΔPSI|>=10%, confidence interval of 95%). Since variations in gene expression levels can affect the accuracy with which AS can be detected and quantified ^30^, we excluded from subsequent analysis all AS events associated with genes that undergo statistically significant changes in gene expression (log2FC > 1 and FDR <0.05) in the same pairwise comparison **(Table S2)**. From this filtering, we obtained 371 DAS events associated with 295 genes **(Table S3)**. Remarkably, only a minority of genes associated with DAS events undergoes changes in gene expression in the same comparison **(Figure S1C)**. In addition, we found that most of the genes that undergo DAS are expressed at high levels (80%), as compared to the distribution of all expressed genes (∼50%) **(Figure S1D)**. Thus, these results suggest that the majority of alterations in AS are independent of changes in gene expression levels of those same genes.

Next, we explored the dynamics of DAS during lung development. We performed K-means clustering of DAS events that occur in non-differentially expressed genes **(Figure 1B)**, specifying the optimal number of clusters to 2, as determined by the average silhouette width method **(Figure S1E)**. Remarkably, clustering of AS events segregated them between embryonic and postnatal stages. Cluster 1 contains AS events whose percent spliced in (PSI) values decrease postnatally; and cluster 2 contains AS events whose PSI values increase postnatally **(Figure 1B-C, Table S5)**, with the majority of the AS changes detected occurring between E18.5 and P5. A complementary analysis using hierarchical clustering of these DAS events identifies this same trend **(Figure S1F)**. This result suggests that adaptation to breathing (embryonic-to-postnatal transition) may involve a comprehensive program of AS changes on a large number of genes, which might fine-tune their activity independent of gene expression.

To understand which pathways are associated with changes in AS during the embryonic-to-postnatal transition, we performed KEGG pathway analysis on the DAS events occurring in non-differentially expressed genes. Enriched terms reveal that AS occurs in genes that are associated with adherens junctions (such as *Afdn, Ctnnd1 and Baiap2*), tight junctions (such as *Magi1, Patj and Amotl1*), and HIPPO signaling pathway (such as *Yap1*, *Tead1* and *Llgl2)* **(Figure 1D and Table S6)**. In sum, our results show that significant AS changes occur in the developing mouse lung during the embryonic-to-postnatal transition, between E18.5 and P5. These changes occur in genes involved in cell-cell adhesion complexes and a signaling pathway known to mediate intercellular communication during lung development.

### In silico analysis identifies RBP candidates for regulation of AS during lung development

The fact that most DAS occur between E18.5 and P5 suggests that these AS events may be regulated by a common regulatory mechanism. AS changes are often driven by changes in the levels of RNA binding proteins (RBPs) ^31^ Thus, we sought to identify RBPs that could regulate the AS changes in lungs. We searched for motif enrichment/depletion on DAS events between E18.5 and P5 occurring in non-differentially expressed genes. We tested 250 nt sequences flanking the splice sites of all regulated splicing events (intronic and exonic) against all RNA-binding proteins in the CISBP-RNA database ^32^ using Matt ^33^. For exon skipping events, we found a significant result (either enrichment or depletion) for 49 motifs, corresponding to 39 RBPs, while for intron retention events we found 64 motifs corresponding to 45 RBPs **(Figure 2A and Table S7)**. We then focused on RBPs that change their expression during development, as it has been shown that these could drive the inclusion/skipping of AS events ^11^. Thus, from the 363 mouse RBPs listed on the CISBP-RNA database, we filtered those that undergo differential gene expression between E18.5 and P5. We identified 47 RBPs fulfilling this condition, 11 of which increased in expression and 36 decreased in expression from E18.5 to P5 **(Figure 2B, Table S8)**. Remarkably, only 3 out of 47 RBPs exhibit both a significant enrichment of their motifs and differential gene expression between E18.5 and P5: *Hnrnpa1* (downregulated), *Cpeb4* (upregulated) and *Elavl2* (downregulated) **(Figure 2C, Table S9)**. Our results show that included exons have more motifs for the upregulated *Cpeb4* and for the downregulated *Elavl2*, while retained introns have more motifs for the downregulated *Hnrnpa1* **(Figure S2A)**. Of note, we could validate the binding of hnRNP A1 to OGT intron 4 and its conservation in human cells using CLIP-seq profiles **(Figure S2B)**, supporting the possibility that the binding events identified may indeed occur in cells. These results position Cpeb4, Elavl2/HuB and hnRNP A1 as strong candidate RBPs for the regulation of the exon skipping and intron retention events detected in the mouse lungs on the embryonic to postnatal transition. Next, we re-analyzed the AS events that are associated with differentially expressed genes and searched for AS events enriched in Cpeb4, Elavl2/HuB and hnRNP A1 binding motifs. Out of 89 AS events, associated with 59 genes, we identified 21 AS events containing one or more of these binding motifs **(Table S10)**, including genes such as *Abr*, *Aspn* and *Vegfa*, which have been previously associated with lung development and/or pathology ^7, 8, 34, 35^. In conclusion, we found a comprehensive and specific AS signature and potential AS regulators involved in the embryonic to postnatal transition.

**Figure 2.**
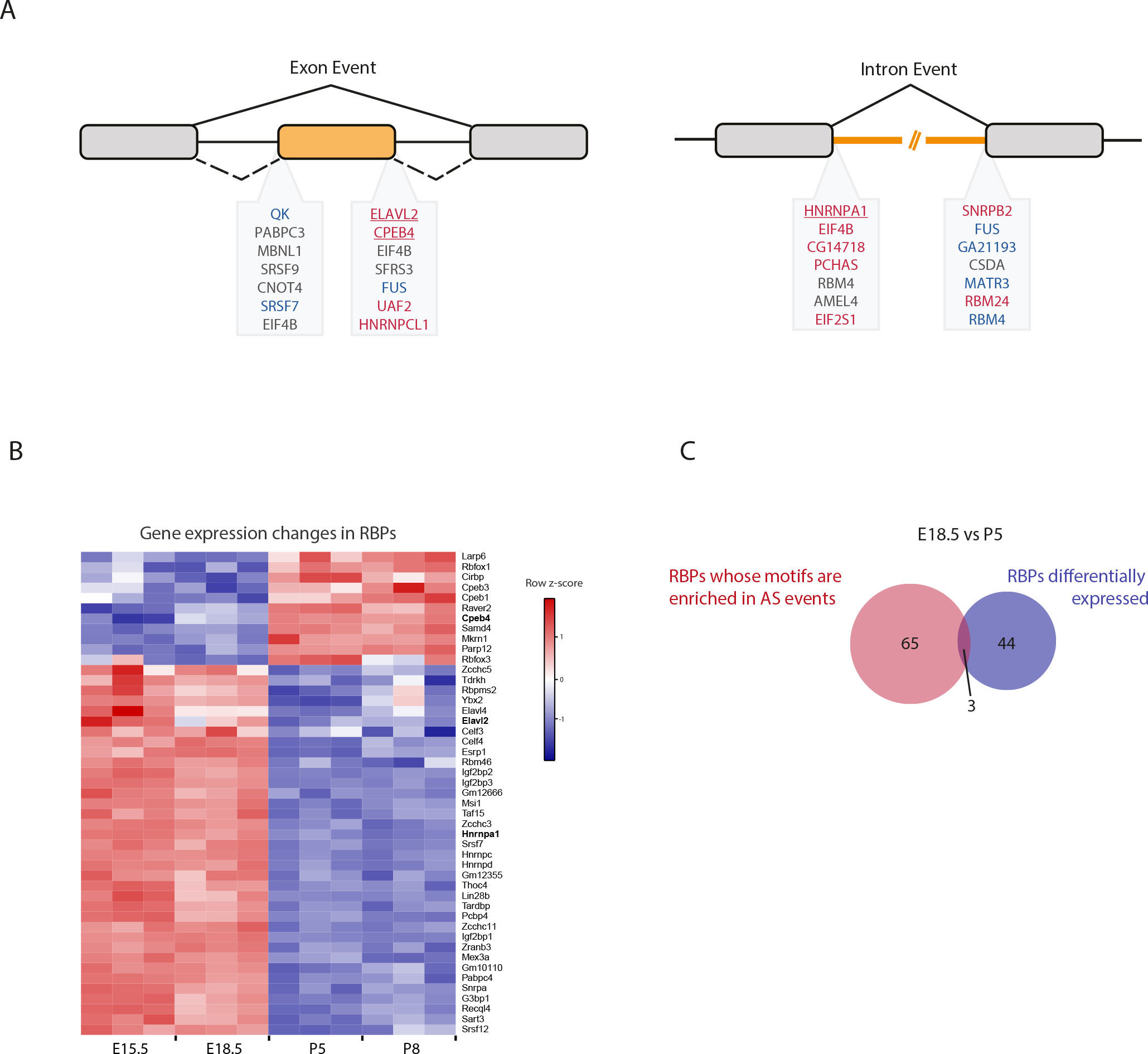
Candidate RBPs for the regulation of DAS between E18.5 and P5. A. RBPs whose motifs are enriched in DAS events occurring in non-differentially expressed genes between E18.5 and P5. Red, enrichment in enhanced vs unregulated. Blue, enrichment in silenced vs unregulated. Gray, depletion in silenced/enhanced vs unregulated. Bold and underlined, RBPs whose motifs are enriched and that undergo differential gene expression between E18.5 and P5. B. Heatmap representing gene expression changes of RBPs that undergo differential gene expression between E18.5 and P5 and whose motifs were enriched in the AS events. Blue and red represent decreased and increased gene expression, respectively, relative to the mean of each gene across the time course (row z-score calculated from log2 (CPMs+1). C. Venn diagram representing the overlap between the RBPs with enriched motifs in DAS events occurring in non-differentially expressed genes between E18.5 and P5 and those that undergo differential gene expression genes between E18.5 and P5

### Analysis of AS identifies a novel *Vegfa* isoform containing intron 5

Next, to further dissect DAS relevant for the transition between embryonic and post-natal time-points, we focused on *Vegfa,* since it is essential for lung development ^7, 8^ by regulating blood vessel formation ^36^. Visual inspection of *Vegfa* RNAseq profile revealed the increased inclusion of *Vegfa* exon 6 and of intron 5 into processed mRNA transcripts from embryonic to postnatal time-points **(Figure 3A)**. While AS of *Vegfa* exon 6 has been previously described ^22^, the inclusion of intron 5 in *Vegfa* mature mRNA species has never been reported before. Thus, we characterized in more detail the existence of intron 5-containing isoforms. To identify which *Vegfa* isoform(s) contain(s) intron 5, we analysed RNA extracted from lungs at P5, a stage at which intron 5 retention was evident **(Figure 3A)**. Specifically, from these RNA samples, we produced cDNA and performed end-point PCR amplification of the intron 5- containing cDNA molecules. For that, we used primer pairs in which one primer anneals with intron 5 and the other anneals with the 5’UTR/exon 1 or with 3’UTR **(Figure 3B)**. The size of the resulting PCR products and the fact that only one band per PCR reaction was obtained **(Figure 3C)**, suggested that the *Vegfa* isoform containing intron 5 also contains all *Vegfa* exons from 1 to 8. We named this isoform *Vegfa i5*. The sequence of this newly identified isoform was further confirmed by Sanger sequencing of the amplified PCR products **(Figure 3B, bottom panel)**.

**Figure 3.**
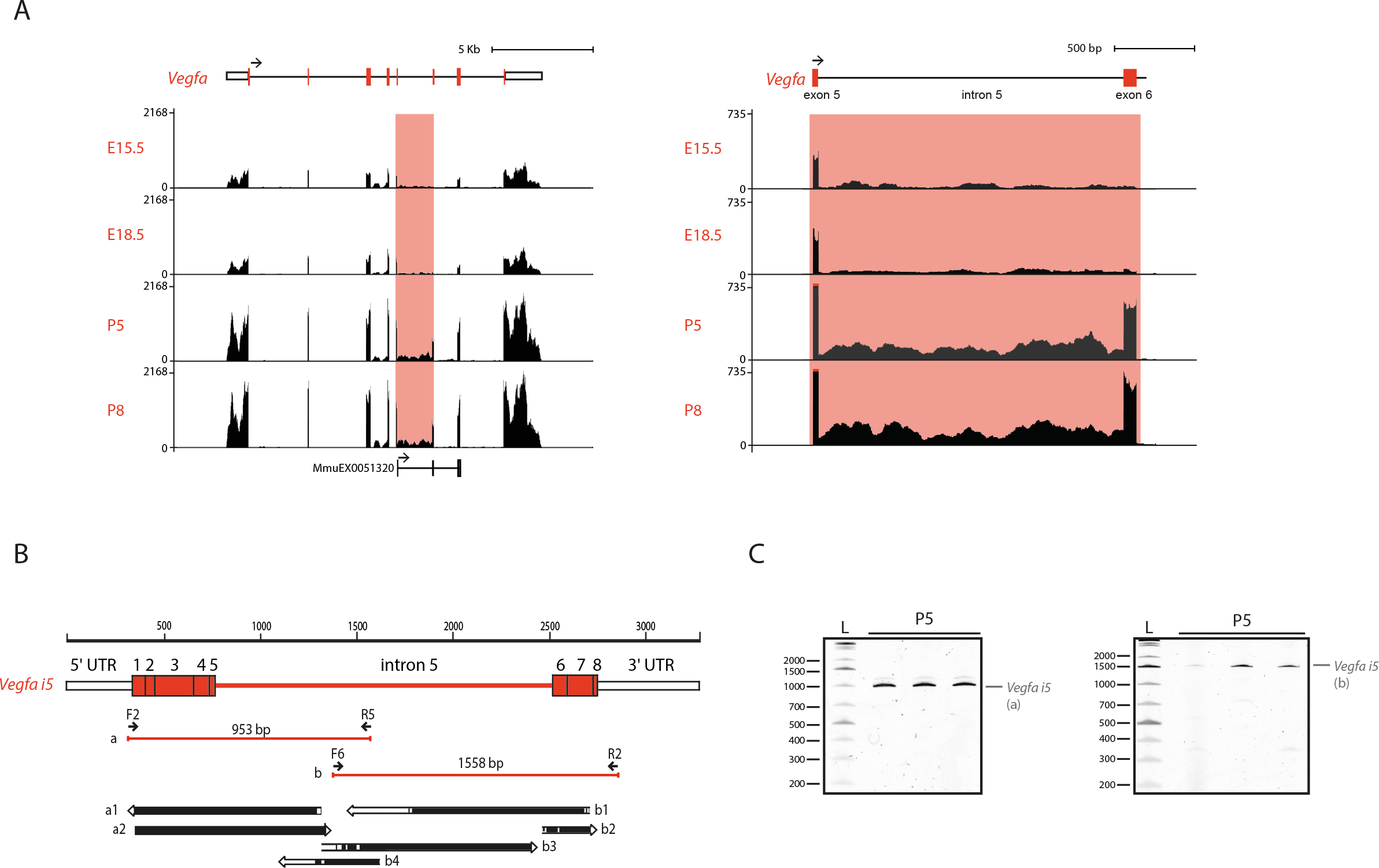
Identification of a novel *Vegfa* isoform – Vegfa *i5*. A. Left: RNAseq profile of *Vegfa* from bulk lung at different developmental time-points. Vast-TOOLS tracks representing previously annotated AS events associated with *Vegfa*. Right: Higher magnification of the region containing intron 5 and exon 6 B. Schematic representation of the identified *Vegfa i5* isoform. Primer pairs used and size of the respective end-point PCR amplicons obtained is indicated. The alignment of sequenced PCR products with the predicted *Vegfa i5* isoform is represented on the bottom panel. Black bars represent full alignment of the sequence, white bars indicate no alignment. Poor sequencing quality at the ends of the sequenced fragments justifies the predominance of non-aligned sequences in these regions. C. TBE-Urea PAGE gels of the *Vegfa i5* amplification products indicated in Figure 3B.

Interestingly, we predicted motif hits for all RBPs in the Catalog of Inferred Sequence Binding Preferences (CISBP-RNA) database along the Vegfa intron 5 sequence and found 19 matches that correspond to differentially expressed genes through development **(Table S11)**. These include the RBPs above identified as candidate regulators of the AS changes, Elavl2/HuB, Cpeb4 and hnRNP A1, suggesting that these may also regulate *Vegfa i5* during mouse lung development. These results revealed a previously unidentified *Vegfa* isoform to be expressed during lung development.

### *Vegfa* isoforms expression changes during lung development

To further identify which *Vegfa* isoforms, in addition to *Vegfa i5*, are expressed during lung development, we analysed the expression changes of the *Vegfa* isoforms previously annotated in ENSEMBL using Kallisto ^37^. Moreover, we manually annotated the newly identified *Vegfa i5* isoform on Kallisto index. Of the annotated isoforms, we found that *Vegfa 120*, *164*, *188* and *i5* are expressed during lung development and that all these isoforms increase in expression from embryonic to postnatal time-points **(Figure 4A)**. This increase is concordant with the increase in total *Vegfa* expression levels in RNAseq during the perinatal period, from E18.5 to P5 **(Figure S3A)**. Remarkably, the isoforms showing the most prominent increase are the ones containing exon 6: *Vegfa 188* and *i5*(absolute fold change from E18.5 to P5 of 7.06 and 4.18, respectively) **(Figure 4A)**. Concordantly, the *Vegfa* Vast DB AS event undergoing changes is the inclusion of exon 6 **(Figure 3B)**.

**Figure 4.**
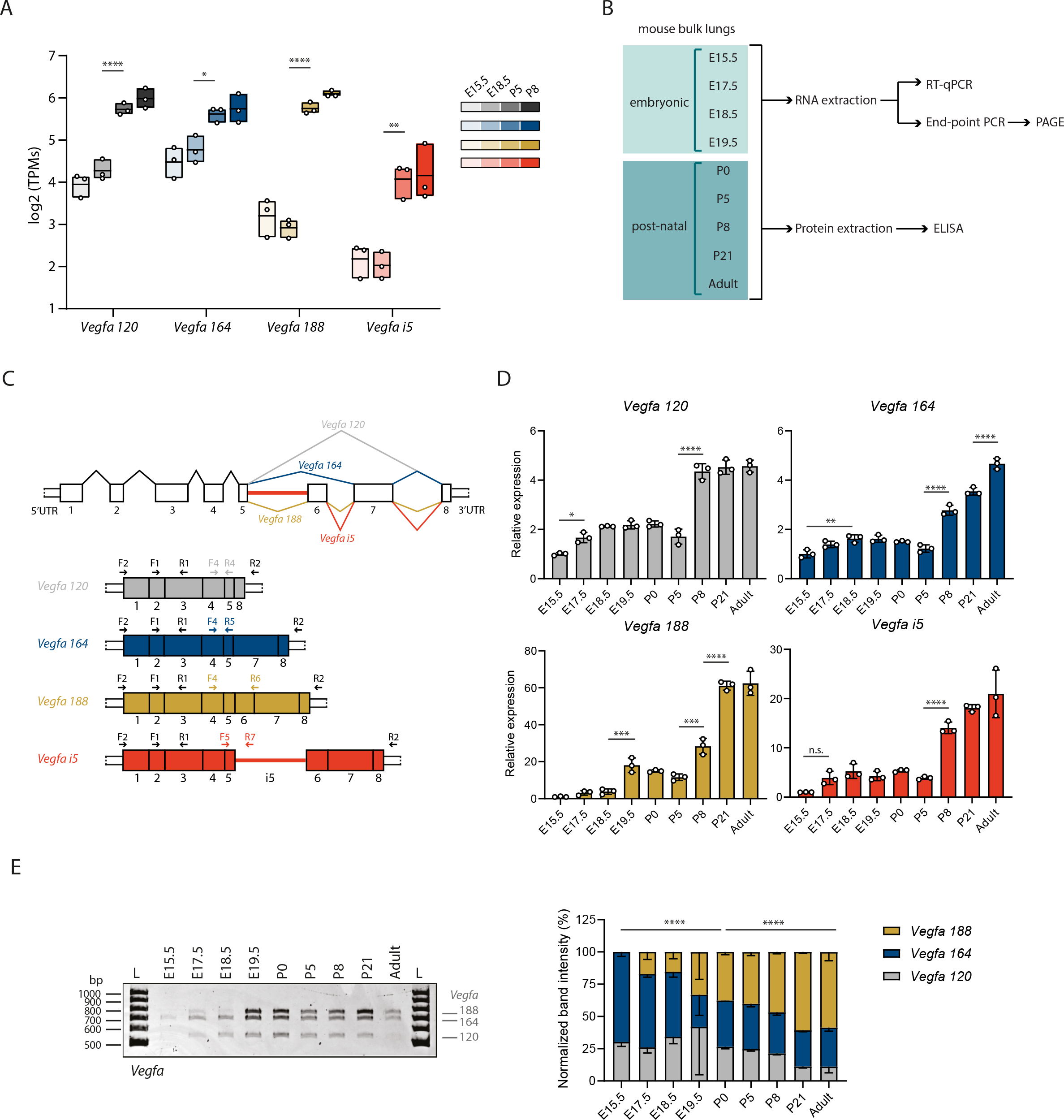
*Vegfa* isoforms are dynamically expressed during lung development. A. Quantification of expression of *Vegfa* AS isoforms at different developmental time-points using Kallisto. TPM, transcripts per million. p-value from one-way ANOVA with Tukey correction for multiple testing. Data represented as mean ± Max/Min. B. Schematic representation of workflow: Bulk lungs at different developmental time-points were collected. RNA or protein were extracted. cDNA was produced from RNA. Expression analysis was performed by RT-qPCR. Analysis of the relative proportions between the expression of different isoforms was performed by end-point PCR and PAGE. Protein levels were quantified by ELISA. C. Schematic representation of the *Vegfa* isoforms analysed in this study. Each exon is represented by a number and *i5* represents intron 5. Primer pairs used to amplify all *Vegfa* isoforms and each isoform individually in RT-qPCR or end-point PCR are indicated in the figure. D. Expression changes of *Vegfa* isoforms from bulk lungs at different developmental time- points analysed by RT-qPCR. N=3 for each time-point. p-value from one-way ANOVA with Tukey correction for multiple testing. E. Left: Representative TBE-Urea PAGE from PCR products obtained from cDNA samples from bulk lungs at different developmental time-points. Fragments were amplified using primer pair FX-RX. Bands represent *Vegfa 188*, *164* and *120* isoforms. Right: Quantification of the normalized relative proportions between the expression levels of *Vegfa* isoforms on bulk lungs at different developmental time-points. N=3 for each time- point. p-value from chi-square test. TBE-Urea PAGE gels used for this quantification is represented in **Figure S5E**.

To characterize the temporal dynamics of AS of *Vegfa* with higher resolution, we collected lungs at E15.5, E17.5, E18.5, E19.5, P0, P5 and P8, as well as two later time-points: P21, at which alveologenesis is still occurring, and adult, at which alveologenesis has already ceased **(Figure 4B)**. The analysis of the expression dynamics of each *Vegfa* isoform at different developmental stages was performed by RT-qPCR. For that, we designed primer pairs that specifically amplify each of the isoforms **(Figure 4C)**. In addition, we designed a pair of primers that amplifies all *Vegfa* isoforms **(Figure 4C**, primers F2 and R2). The specificity of these primer pairs on the RT-qPCR was validated by the amplification of a single PCR product of the predicted size for each isoform **(Figure S3C)**. This result also excluded the eventual occurrence of spurious amplification of genomic DNA.

In accordance with the RNAseq results, RT-qPCR analysis revealed a progressive increase in total *Vegfa* mRNA levels from E15.5 to the adult stage **(Figure S3D)** which is associated with an increase of the VEGFA protein levels, as assessed through ELISA analysis of whole lung protein lysates **(Figure S3E)**. From the *Vegfa* isoforms, we found that there is a significant increase in *Vegfa 120* and *164* from E15.5 to P8 (fold change of 4.35 and 2.78, respectively) **(Figure 4D)**. The expression levels of these isoforms remain high at P21 and adult lungs. On the other hand, *Vegfa 188* levels sharply increase at E19.5 (fold change of 18.09 between E15.5 and E19.5), just before birth and then increase even further at P8, P21 and adult (fold change of 28.38, 61.13 and 62.46 between E15.5 and P8, E15.5 and P21, and E15.5 and Adult, respectively) **(Figure 4D)**. *Vegfa i5* increases from E15.5 to E17.5 (fold change of 3.89), and further increases at P8 (fold change of 14.06 between E15.5 and P8), remaining high at P21 and adult **(Figure 4D)**. Importantly, the increase in *Vegfa 188* and *i5* is much higher than that of *Vegfa 120* and *164* (fold change of 28.38, 14.06 compared to 4.35, 2.78 between E15.5 and P8, respectively). These observations suggest that the relative proportion between *Vegfa* isoforms changes during lung development. Although RT-qPCR allows the accurate quantification of each isoform, it is not the most suitable method to compare relative changes in expression levels between multiple isoforms.

To be able to perform this analysis, we performed end-point PCR followed by polyacrylamide gel electrophoresis (PAGE). For that, we used a single primer pair that hybridizes at *Vegfa* 5’ and 3’ UTRs and that amplifies the *Vegfa* isoforms *120*, *164* and *188* **(Figure 4C**, primers F1 and R1). The amplification of these 3 isoforms was detectable by the presence of three distinct bands with the size corresponding to each of these isoforms **(Figure 4E)**. Although this pair of primers should theoretically be able to amplify the *Vegfa i5* isoform as well, it does not. This happens probably due to its larger size when compared to the other isoforms (2511 bp vs 571, 703 and 775 bp) and to its lower level of expression **(Figure 4A)**, which makes it more difficult to be amplified when in competition with the lower size and abundant isoforms. Nevertheless, we could use this technique to evaluate changes in the proportions between the expression of isoforms *Vegfa 120*, *164* and *188*. This analysis revealed that *Vegfa 164* is the isoform more predominantly expressed in bulk lungs at the embryonic time-points, followed by *Vegfa 120*, with only a minor contribution from *Vegfa 188*. From E17.5 onwards, the proportion of *Vegfa 188* gradually increases, reaching a maximum of 70% in the adult. Reciprocally, the relative proportions of both *120* and *164* decrease **(Figure 4E, S3F)**. These results are concordant with previously published results obtained using RNA protection assays from RNA extracted from mouse lungs during development ^16^.

In sum, we found that all the detected *Vegfa* isoforms start to progressively increase before birth and further increase after birth, which coincides with an overall increase in total *Vegfa* levels. However, they increase at distinct rates during lung development: *Vegfa 188* and *Vegfa i5* isoforms undergo a marked differential increase, when compared to that of *120* and *164*. This demonstrates the occurrence of *Vegfa* AS during lung development towards the expression of the exon 6-containing isoforms. Remarkably, our fine-grained analysis shows an increase in the relative proportion of *Vegfa 188* starting before birth. These observations suggest that *Vegfa* AS coincides with a developmental adaptation to birth.

### *Vegfa* isoforms are differentially expressed between endothelial and epithelial populations during lung development

Our results showed that *Vegfa* undergoes AS changes during lung development. However, it was unclear which cell types express which *Vegfa* isoforms and what is their expression dynamics within the different cell types. Previously, *Vegfa* expression was documented in ECs, AT1 and AT2 cells by *in situ* hybridization ^16, 17, 24^. More recently, genetic LacZ reporters and scRNAseq analyses have reported the expression of *Vegfa* only in AT1 and ECs ^8, 14, 38, 39^. Yet, these studies have only examined global levels of *Vegfa* transcripts. To unravel which cells express the different *Vegfa* isoforms, we isolated various lung cell types at different time- points of development and assessed how *Vegfa* isoforms expression varies within each cell type **(Figure 5A)**. We examined lungs at E15.5, E17.5, E18.5, E19.5, P0, P5, P8, P21 and adult, in accordance with our analysis in bulk lungs. To isolate the different lung cell types, we dissociated the lung tissue into a single-cell suspension and performed fluorescence-activated cell sorting (FACS) using antibodies for cell type-specific markers. The combination of these markers allowed the isolation of cell populations enriched for endothelial cells (CD31 single positive (SP): CD31^+^, EpCAM^-^, CD45^-^), epithelial cells (EpCAM SP: CD31^-^, EpCAM^+^, CD45^-^), immune cells (CD45 SP: CD31^-^, EpCAM^-^, CD45^+^) and mesenchymal cells, such as alveolar myofibroblasts and pericytes (triple negative (TN): CD31^-^, EpCAM^-^, CD45^-^) **(Figure 5B)**. The analysis of gene expression changes was performed by RNA extraction from the isolated cell populations followed by RT-qPCR or end-point PCR **(Figure 5A)**.

**Figure 5.**
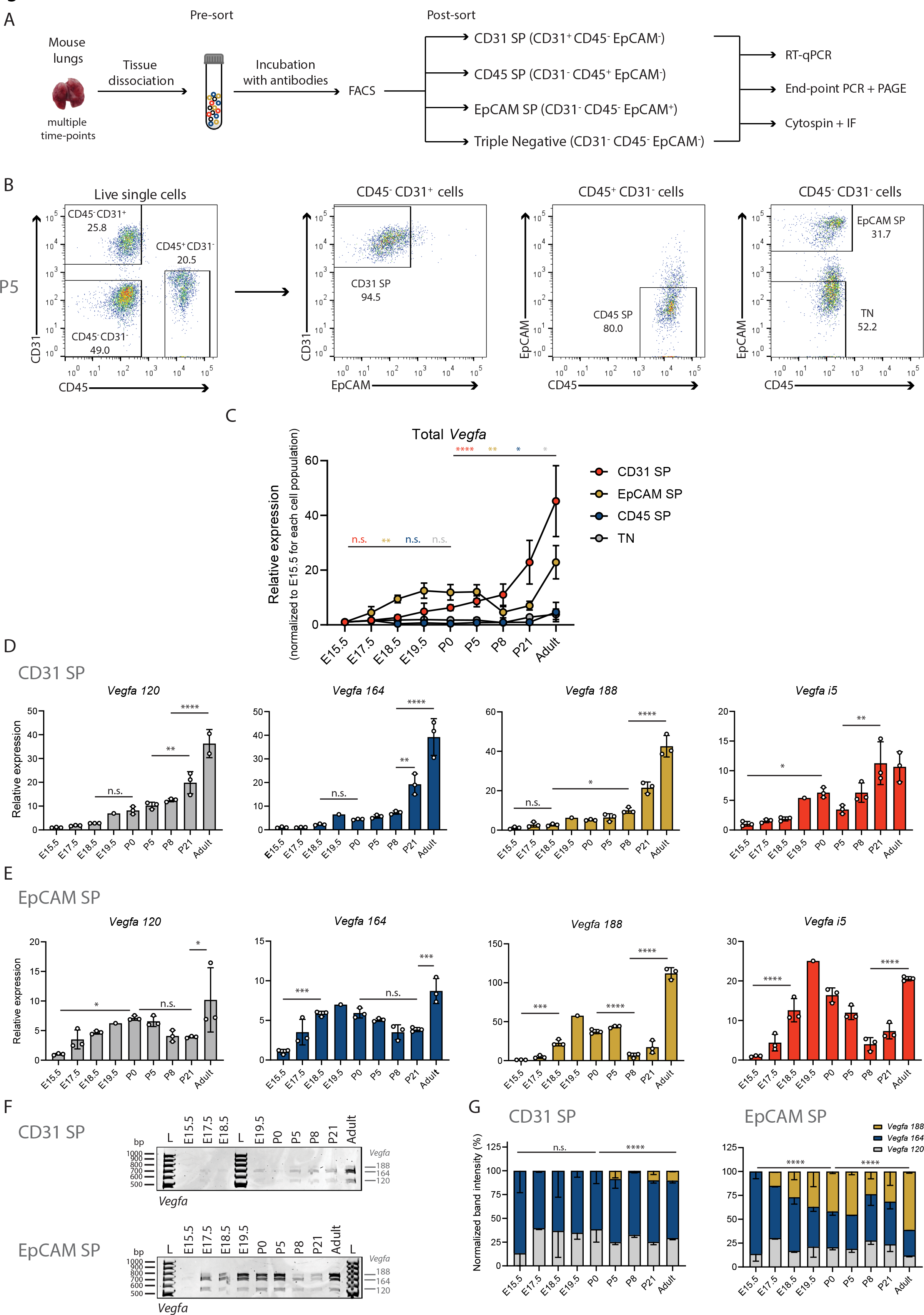
*Vegfa* isoforms are differentially expressed between CD31 SP and EpCAM SP populations during lung development. A. Schematic representation of workflow: Cell suspensions from lungs at different developmental time-points were collected (pre-sort sample). Several lung cell populations were isolated by fluorescent-activated cell sorting (FACS). Samples were analysed for the expression of indicated surface markers (CD45, CD31, EpCAM). CD31 SP populations are enriched for endothelial cells, CD45 populations are enriched for immune cells, EpCAM SP populations are enriched for epithelial cells, Triple negative are enriched for mesenchymal cells. RNA was extracted from isolated cell populations and cDNA produced. Expression analysis was performed by RT-qPCR. Analysis of the relative proportions between the expression of different isoforms was performed by end-point PCR and PAGE. Alternatively, isolated cell populations were processed by cytospin and immunofluorescence (IF) was performed. B. Representative FACS analysis plot from lung cell suspensions at P5. Dot plots represent cell populations from indicated gates. Live cells were gated using LiveDead Fixable Viability Dye. Percentages from indicated populations are represented. C. Expression changes of total *Vegfa* from sorted cell populations at different developmental time-points analysed by RT-qPCR. Expression values normalized to E15.5 for each cell population. N=3 for each time-point. p-value from one-way ANOVA with Tukey correction for multiple testing. D. Expression changes of *Vegfa* isoforms from CD31 SP cell population at different developmental time-points analysed by RT-qPCR. N=3 for each time-point except for E19.5. N=1 for E19.5. p-value from one-way ANOVA with Tukey correction for multiple testing. E. Expression changes of *Vegfa* isoforms from EpCAM SP cell population at different developmental time-points analysed by RT-qPCR. N=3 for each time-point except for E19.5. N=1 for E19.5. p-value from one-way ANOVA with Tukey correction for multiple testing. F. Representative TBE-Urea PAGE from PCR products obtained from cDNA samples from CD31 SP and EpCAM SP cell population at different developmental time-points. G: Quantification of the normalized relative proportions between the expression levels of *Vegfa* isoforms on CD31 SP and EpCAM SP cell population at different developmental time- points. N=3 for each time-point. p-value from chi-square test. TBE-Urea PAGE gels used for this quantification are represented in **Figure S6B-C**.

We first evaluated the quality of the isolation of the different cell populations. For that, we examined the expression of cell type-specific markers in the different cell populations collected at P5 by RT-qPCR. We analysed the mRNA levels of *Pecam1* (CD31) and *Cdh5* as pan-endothelial-specific markers, of *Prox1* as a marker for lymphatic ECs, of *Epcam* and *Cdh1* as pan-epithelial-specific markers, and of *Cd45* as a pan-immune cell marker. We identified that only CD31 SP population expresses *Pecam1* (CD31) and *Cdh5* **(Figure S4A)**. However, this population also expresses high levels of *Prox1* **(Figure S4A)**, suggesting that CD31 SP contains a mixture of blood and lymphatic ECs. We also showed that only EpCAM SP population expresses *Epcam* and *Cdh1*, and that only CD45 SP expresses *Cd45*, while TN cells do not express any of these markers **(Figure S4A)**. In addition, by analyzing the mRNA levels of *Aqp5*, *Sftpc* and *Foxj1*, markers of epithelial alveolar AT1, AT2 and ciliated cells, respectively, we demonstrated that the EpCAM SP population is composed by a mixture of these lung epithelial subtypes **(Figure S4A)**.

To estimate the contribution of each endothelial and epithelial subtypes in CD31 SP and EpCAM SP populations, we analysed the expression of cell type-specific markers at the protein level in single cell suspensions before sorting (pre-sort) and after sorting using cytospin preparation ^40^ and immunostaining. We observed that CD31 SP population is highly enriched for CD31-positive cells, as compared to the pre-sort sample. Of these, about 60% are also positive for ERG, a marker of blood endothelial cells **(Figure S4B)**. The remaining fraction of CD31-positive cells is likely composed of lymphatic ECs, as suggested by the enrichment in expression of the lymphatic ECs marker *Prox1* in this population **(Figure S4A)**. EpCAM SP population is enriched for AQP5-positive and SFTPC-positive cells (21.2% and 79.2% on average, respectively), demonstrating that this population is enriched for both epithelial AT1 and AT2 cells **(Figure S4C-D)**. Altogether, these results validate the quality of our method for isolation of different lung cell types.

We then analysed the expression changes of each *Vegfa* isoform in each cell population along lung development by RT-qPCR. Total *Vegfa* expression is highest in EpCAM SP in all time- points analysed, followed by CD31 SP population **(Figure S5A)**. In addition, we found that CD31 SP and EpCAM SP populations reveal the highest fold changes in total *Vegfa* expression during development, when compared to CD45 SP and TN populations (fold change between E15.5 and adult of 45.21, 22.90, 4.70, and 3.87, respectively) **(Figure 5C)**. Therefore, to characterize *Vegfa* AS, we focused our analysis in EpCAM SP and CD31 SP populations We found that in the CD31 SP population *Vegfa 120*, *164*, *188* and *i5* steadily increase during lung development from E18.5 to adult **(Figure 5D)**. The analysis of *Vegfa* isoforms relative proportions by end-point PCR and PAGE revealed that *Vegfa 164* is the most abundant isoform and that there was no significant change in the proportion between the *Vegfa* isoforms during lung development in CD31 SP population **(Figure 5F-G, S5B)**. These results suggest that AS of *Vegfa* does not significantly change in the EC population.

In EpCAM SP population, *Vegfa 120* and *164* increase before birth from E15.5 onwards, peaking at P0, after which decrease until P21. *Vegfa 188* and *i5* increase from E17.5 until P5 **(Figure 5E)**. In the adult, *Vegfa 120*, *164* and *188* exhibit the highest levels both in EpCAM SP and CD31 SP populations **(Figure 5D-E)**.

Analysis of the relative proportions of *Vegfa* isoforms in EpCAM SP population through end- point PCR and PAGE showed that there is a prominent increase in the relative proportion of *Vegfa 188*in epithelial cells throughout lung development **(Figure 5F-G, S5C)**. While in embryonic time-points *Vegfa 164* and *Vegfa 120* are the predominant isoforms, the proportion of *Vegfa 188* increases progressively throughout development and, at postnatal time-points, it becomes the most abundant isoform expressed in EpCAM SP cells **(Figure 5G, S5C)**. These results suggest that the signature of AS for *Vegfa* that we have first detected on bulk lungs is associated with changes occurring in the epithelial lineage.

In sum, our results suggest that the endothelial and epithelial lineages express the highest *Vegfa* levels and that *Vegfa* expression increases in both during lung development. While in ECs, *Vegfa 164* is always the predominant isoform, in epithelial cells there is a marked increase in the proportion of *Vegfa 188*, from E17.5 onwards. These results suggest a cell type-specific AS of *Vegfa* in epithelial cells during lung development at the perinatal period.

### *Vegfa* undergoes AS in epithelial AT1 cells during lung development

The EpCAM SP population is composed of multiple subtypes such as epithelial alveolar AT1, AT2 and epithelial bronchiolar ciliated cells **(Figure S4A, C, D)**. To dissect the contribution of each epithelial cell subtype for the expression of *Vegfa* isoforms, we further subdivided EpCAM-positive population into AT1- and AT2-enriched subpopulations by FACS in P5 lungs **(Figure 6A)**, a stage at which *Vegfa* expression in the epithelial lineage is high **(Figure S5A)**. For that, we used the marker major histocompatibility complex class II (MHC II) which, in combination with EpCAM, has been previously shown to discriminate between AT1 (EpCAM^low^ MHC II^-^), AT2 (EpCAM^high^ MHC II^+^), and ciliated cells (EpCAM^high^ MHC II^-^) ^41^. We could identify within the EpCAM-positive cell population the presence EpCAM^low^ MHC II^-^ and EpCAM^high^ MHC II^+^ subpopulations putatively corresponding to AT1 and AT2, respectively **(Figure 6B)**. However, we detected very few EpCAM^high^ MHC II^-^ cells, putative ciliated cells, and, therefore, this subpopulation was not further analysed **(Figure 6B)**.

**Figure 6.**
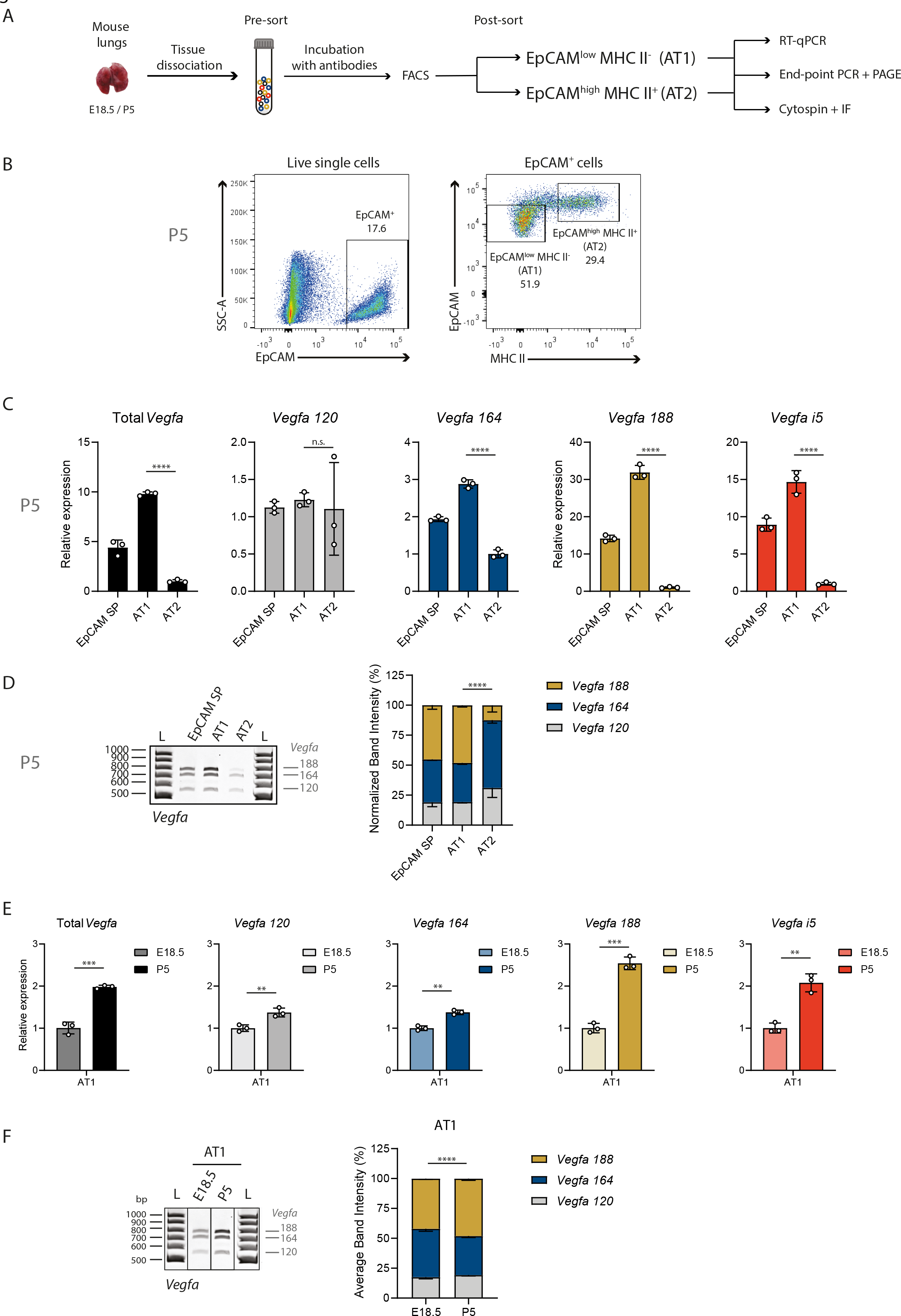
*Vegfa* undergoes alternative splicing towards *Vegfa 188* in epithelial AT1 during lung development. A. Schematic representation of workflow: Cell suspensions from lungs at E18.5 and P5 were collected (pre-sort samples). Several lung cell populations were isolated by fluorescent- activated cell sorting. Samples were analysed for the expression of indicated surface markers (EpCAM, MHC II). EpCAM^low^ MHC II^-^ populations are enriched for AT1 cells, EpCAM^high^ MHC II^+^ populations are enriched for AT2 cells. RNA was extracted from isolated cell populations and cDNA produced. Expression analysis was performed by RT-qPCR. Analysis of the relative proportions between the expression of different isoforms was performed by end-point PCR and PAGE. Alternatively, isolated cell populations were processed by cytospin and immunofluorescence (IF) was performed. B. Representative FACS analysis plot from lung cell suspensions at P5. Dot plots represent cell populations from indicated gates. Live cells were gated using LiveDead Fixable Viability Dye. Percentages from indicated populations are represented. C. Expression changes of *Vegfa* isoforms from EpCAM SP, AT1 and AT2 cell population at P5 were analysed by RT-qPCR. N=3 for each time-point. p-value from unpaired t-test. D. Left: Representative TBE-Urea PAGE from PCR products obtained from cDNA samples from EpCAM SP, AT1 and AT2 cell populations at P5. Right: Quantification of the normalized relative proportions between the expression levels of *Vegfa* isoforms on EpCAM SP, AT1 and AT2 cell populations at P5. N=3 for each time-point. p- value from chi-square test. TBE-Urea PAGE gels used for this quantification are represented in Figure S3F and S4E. E. Expression changes of *Vegfa* isoforms from AT1 cell population at E18.5 and P5 were analysed by RT-qPCR. N=3 for each time-point. p-value from unpaired t-test. F. Left: Representative TBE-Urea PAGE from PCR products obtained from cDNA samples from AT1 cell populations at E18.5 and P5. Right: Quantification of the normalized relative proportions between the expression levels of *Vegfa* isoforms on AT1 cell populations at E18.5 and P5. N=3 for each time-point. p-value from chi-square test. TBE-Urea PAGE gels used for this quantification are represented in Figure S4E.

mRNA analysis of the selected subpopulations by RT-qPCR revealed that none of these two subpopulations expresses *Pecam1* (CD31) nor *Cd45*, while both express *Epcam* mRNA **(Figure S6A)**. In addition, by RT-qPCR, we found that EpCAM^low^ MHC II^-^ cells express high levels of *Aqp5* mRNA and low levels of *Sftpc* mRNA, while EpCAM^high^ MHC II^+^ cells express high levels of *Sftpc* mRNA and low levels of *Aqp5* mRNA **(Figure S6B)**, suggesting that indeed they are enriched for AT1 and AT2 cells, respectively. None of these two subpopulations is enriched for a marker of epithelial bronchial ciliated cells *Foxj1* mRNA, as compared to EpCAM SP population **(Figure S6B)**, suggesting that this cell type was not collected in this analysis.

Complementarily, by cytospin and immunofluorescence, we found that 54.5% of EpCAM^low^ MHC II^-^ cells are AQP5-positive cells, while 87.8% of EpCAM^high^ MHC II^+^ cells are SFTPC-positive cells **(Figure S6C-D)**. Thus, this analysis revealed that this FACS gating strategy enables the isolation of subpopulations enriched for epithelial AT1 and AT2 cells.

Analysis of *Vegfa* expression levels in AT1- and AT2-enriched subpopulations through RT-qPCR revealed that *Vegfa* isoforms expression is markedly higher in AT1 cells than in AT2 **cells (Figure 6C)**, in agreement with recent scRNAseq results ^8, 39, 42^. While there is only a moderate increase in the expression of *Vegfa 164*, the levels of expression of *Vegfa 188* and *Vegfa i5* are markedly higher in AT1 than in AT2 (fold change of 2.87, 31.60 and 14.53, respectively) **(Figure 6C)**. The levels of *Vegfa 120* are not significantly distinct between both subpopulations. Through end-point PCR and PAGE, we found that *Vegfa 188* is the isoform whose expression is predominant in AT1 cells at P5 **(Figure 6D, S6E)**. Our results suggest that the expression of *Vegfa 188* in EpCAM SP population is due to its expression in AT1 cells.

*Vegfa 188* starts to be expressed in EpCAM SP population at E17.5 **(Figure 5E)**. Interestingly, it is just before this time-point that AT1 cells start differentiating from AT1/AT2 progenitor cells ^3, 15, 39^. This led us to question if the increase in *Vegfa 188* proportion in EpCAM SP population is solely due to an increase in the fraction of AT1 within EpCAM SP population during development, as we detected by FACS **(Figure S6F)**, or if the predominance of *Vegfa 188* within AT1 cells also increases during development. To disentangle between these two possibilities, we analysed *Vegfa* isoforms expression in AT1 cells at E18.5, a time-point just after their specification, and compared it to that in AT1 cells at P5. By RT-qPCR analysis, we found that *Vegfa* isoforms expression increase in AT1 cells from E18.5 to P5. This increase is mild for *Vegfa 120* and *164*, but it is higher for *Vegfa 188* and *Vegfa i5* (fold change of 1.37, 1.38, 2.54 and 2.08, respectively) **(Figure 6E)**. Accordingly, although at E18.5 *Vegfa 188* proportion is already 42% of the total, its proportion further increases at P5 to 49% **(Figure 6F, S6E)**. Altogether, our results suggest that *Vegfa* undergoes AS changes in epithelial AT1 cells during lung development.

## Discussion

Lung alveolar formation starts during late embryonic development and continues postnatally during the first weeks after birth. The distinct cell types that compose the developing alveoli experience dramatic changes at birth. Oxygen tension, mechanical stretch due to respiratory movements and blood flow rates dramatically increase. However, how the distinct lung cell types adapt to such dramatic changes is still not fully understood.

AS is a wide phenomenon driving developmental changes. However, the regulation of AS during lung development at a genome-wide level had not been explored before. This work represents the first comprehensive analysis of the precise dynamics of AS during lung development at a genome-wide scale. Our analysis uncovered the occurrence of AS in the mouse lung during the embryonic-to-postnatal transition. We identified numerous genes that undergo AS at this transition, creating two clusters that exhibit distinct kinetics between embryonic and postnatal stages. We identified AS changes in key cell-cell adhesion complexes and signaling pathways known to regulate intercellular communication between epithelial, endothelial and other lung cell types during lung development: adherens and tight junctions, and VEGF and Hippo signaling pathways ^7, 8, 49, 50^. Overall, this raises the hypothesis that regulation of AS can be a means of modulating intercellular communication in lungs towards functional respiration. Remarkably, the number of AS changes in genes that do not undergo gene expression changes was considerably higher than in genes that undergo gene expression changes. Thus, it is tempting to speculate that the program of adaptation to birth through AS is largely independent of the gene expression program operating at this stage. Therefore, we postulate that further investigation into the mechanisms governing AS in lung could be relevant for development and pathology. Supporting this vision, several studies have already associated AS and different lung conditions, such as lung cancer, idiopathic pulmonary fibrosis, and chronic obstructive pulmonary disease ^43, 44^.

Through bioinformatics analysis, we identified Cpeb4, Elavl2 and hnRNP A1 as RPBs potentially regulating DAS in mouse lungs. Cpeb4 regulates polyadenylation of the 3’UTR of mRNA transcripts and thus, the stability and translational output and has previously been shown to bind to and regulate *Vegfa* ^45^. hnRNP A1 has been shown to bind to intronic or exonic splice silencers to regulate splicing of alternative exons ^11^. Elavl2/HuB belongs to the family of Hu proteins that have been implicated in the regulation of AS and alternative polyadenylation ^46^. *Cpeb4* or *Elavl2* knock out mice are viable ^47, 48^, thus suggesting a mild effect in lung development, whilst *Hnrnpa1* knock out mice die perinatally with cardiac defects but analysis of lungs was not reported ^48^. Regardless, further investigation on the biological relevance of each of these factors, and combinatorial effects, is warranted to validate their roles in lung development.

We have further explored AS of *Vegfa* gene. Although changes in the relative proportion of *Vegfa* isoforms have previously been described to occur during development ^16–18^, it was not known which cell types in the lung express the different *Vegfa* isoforms, nor if the switch in *Vegfa* AS occurs in a tissue or cell type–specific manner. We identified that *Vegfa* transcript levels increase both in ECs and in AT1 cells during alveolar development. However, while in the embryonic period, *Vegfa 164* is the predominant isoform expressed in both cell populations, in the postnatal period, *Vegfa 188* becomes predominant specifically in AT1 cells. Our results suggest the occurrence of a cell type-specific AS of *Vegfa* in AT1 cells during late lung development. Interestingly, the sharp increase in *Vegfa 188* expression from E18.5 to E19.5 suggests that epithelial *Vegfa 188* could be associated with a pre-birth adaptation of lungs to the postnatal life. It will be interesting to explore how such cell-type specificity of AS is encoded. Specifically, which splicing regulators control this AS event and to explore if other transcripts undergo AS changes within this developmental time interval of lung development in this cell population.

AT1-derived *Vegfa* expression is important for the specification of Car4-positive ECs (also known as aCap or aerocytes), a specialized alveolar EC subtype, ^8, 38^. Remarkably, the specification of this cell type seems to occur just before birth, around E19.5 ^8^. It is, however, unclear which *Vegfa* isoform(s) drive(s) this effect. We found a significant increase in *Vegfa 188* expression from E17.5 to E19.5 in AT1 cells, reaching around 50% of total *Vegfa* transcripts. This observation led us to hypothesize that *Vegfa 188* could specifically be the isoform driving Car4-positive ECs specification in developing lung alveoli and, thus, AS could represent a means of regulating intercellular communication.

We also found a marked increase in *Vegfa* expression during lung development in ECs, in which *Vegfa 164* is the predominant isoform in all time-points analysed. However, it remains to be determined what is the role of EC-derived *Vegfa* both during development and at the adult stage. Previous results from different authors on the effects in the lung of EC-specific *Vegfa* deletion are conflicting. On one hand, Lee et al. reported that non-inducible *Vegfa* EC-specific knock out resulted in premature death and, in the case of surviving mice, these exhibited lungs with increased chronic inflammation and fibrosis, EC rupture, and collapsed lumen ^51^. On the other hand, Ellis et al. reported that the EC-specific deletion of *Vegfa* at P3 elicits no effects in lung alveolar morphology at P9 ^8^. The reasons for this discrepancy are unclear. It is possible that the *Vegfa* deletion in ECs is compensated by the endogenous or ectopic expression of *Vegfa* in AT1 cells.

Another intriguing observation is that *Vegfa* expression in alveolar AT1 and ECs keeps increasing after birth. In many tissues, the formation of new blood vessels usually occurs in response to hypoxia. Under low oxygen tension, hypoxia-inducible factors (HIFs) are protected from proteolytic degradation, become stable, and activate the expression of a myriad of transcriptional targets, among which is *Vegfa* ^52^. Yet, after birth, lung alveoli become filled with oxygen-rich air. This suggests that *Vegfa* expression in the postnatal lungs must be independent of hypoxia. Understanding the mechanisms that allow the uncoupling between hypoxia and *Vegfa* levels in postnatal lungs deserves attention in the future.

In addition to the previously described *Vegfa* isoforms, we have identified a novel *Vegfa* isoform that has undergone splicing of all introns except of intron 5, to which we called *Vegfa i5*. This isoform is expressed during lung development both in ECs and AT1 cells. Intron retention is a widespread phenomenon shown to have a functional impact in development, physiology, and disease ^53–55^. Isoforms containing retained introns may have multiple fates: to encode functionally distinct alternative protein products, to originate prematurely truncated proteins that are targeted for degradation, or having their mRNAs targeted for degradation through nonsense-mediated decay ^53–55^. However, if this mRNA species was rapidly eliminated after its production, the probability of detecting it through RNAseq would very low. Alternatively, the excision of retained introns from pre-spliced mRNAs has also been shown to allow a rapid production of the mature mRNA in response to extracellular stimuli ^54, 56^. It will be relevant to identify how the generation of this novel *Vegfa* mRNA isoform is controlled, and to address its functional significance during lung development. In addition, it has been described that gene expression changes and RNA polymerase II elongation rate may be coupled with changes in AS ^57^. Since we found that both *Vegfa* gene expression and its AS pattern change during lung development, it will be interesting to address if the occurrence of these events on *Vegfa* is interdependent.

In sum, our work contributes for a better understanding of the mechanisms driving alveologenesis and sets the ground for the study of the role of AS dynamics during lung development.

## Acknowledgements

Mariana Ferreira, Marie Bordone, Nuno Agostinho and Nuno Barbosa Morais (Disease Transcriptomics Lab, iMM) for input on bioinformatics analysis.

Pedro Papotto, Karine Serre (Immuno-Biology & Immuno-Oncology Lab, iMM), Idálio Viegas (Biology of Parasitism Lab, iMM), Isabel Alcobia (Institute of Histology and Developmental Biology, FMUL), Debanjan Mukherjee, Vanessa Zuzarte-Luís (Biology & Physiology of Malaria, iMM), and Mahak Singhal (Vascular Oncology and Metastasis Division, DKFZ), for input on experimental protocols.

Luís Oliveira (Cell Architecture Lab, iMM) for input on microscopy image analysis. Flow Cytometry, Bioimaging and Rodent facilities at iMM for technical support.

All members of the Vascular Morphogenesis lab at iMM for discussions, helpful input and for carefully reviewing the manuscript.

Nuno Barbosa Morais for carefully reading this manuscript and providing helpful and critical feedback.

## Author contribution

According to CRediT – Contributor Roles Taxonomy: Conceptualization - MFF, CGF, FFV, CAF

Data curation - MFF, CGF, TB, AARFR, FFV, PC, ARG

Formal Analysis - MFF, CGF, TB, AARFR, FFV, PC, ARG

Funding acquisition – FFV, CAF

Investigation - MFF, CGF, TB, AARFR, FFV, PC, ARG Methodology – FFV, MFF, CGF, CAF, PC, ARG

Project administration – FFV, CAF Resources - MFF, CGF, TB, AARFR, FFV Software - CGF, AARFR, FFV, PC, ARG

Supervision – FFV, CAF

Validation – MFF, CGF, FFV, PC, ARG Visualization – MFF, CGF, FFV, PC, ARG Writing – original draft – FFV, MFF

Writing – review & editing - MFF, CGF, TB, AARFR, FFV, CAF, PC, ARG

## Ethics

Animal experimentation: Mice were maintained at the Instituto de Medicina Molecular under standard husbandry conditions and under national regulations, under the license AWB_2015_11_CAF_Polaridade / ex vivo_surplus_not of use.

## Supplemental figure legends

**Figure S1.**
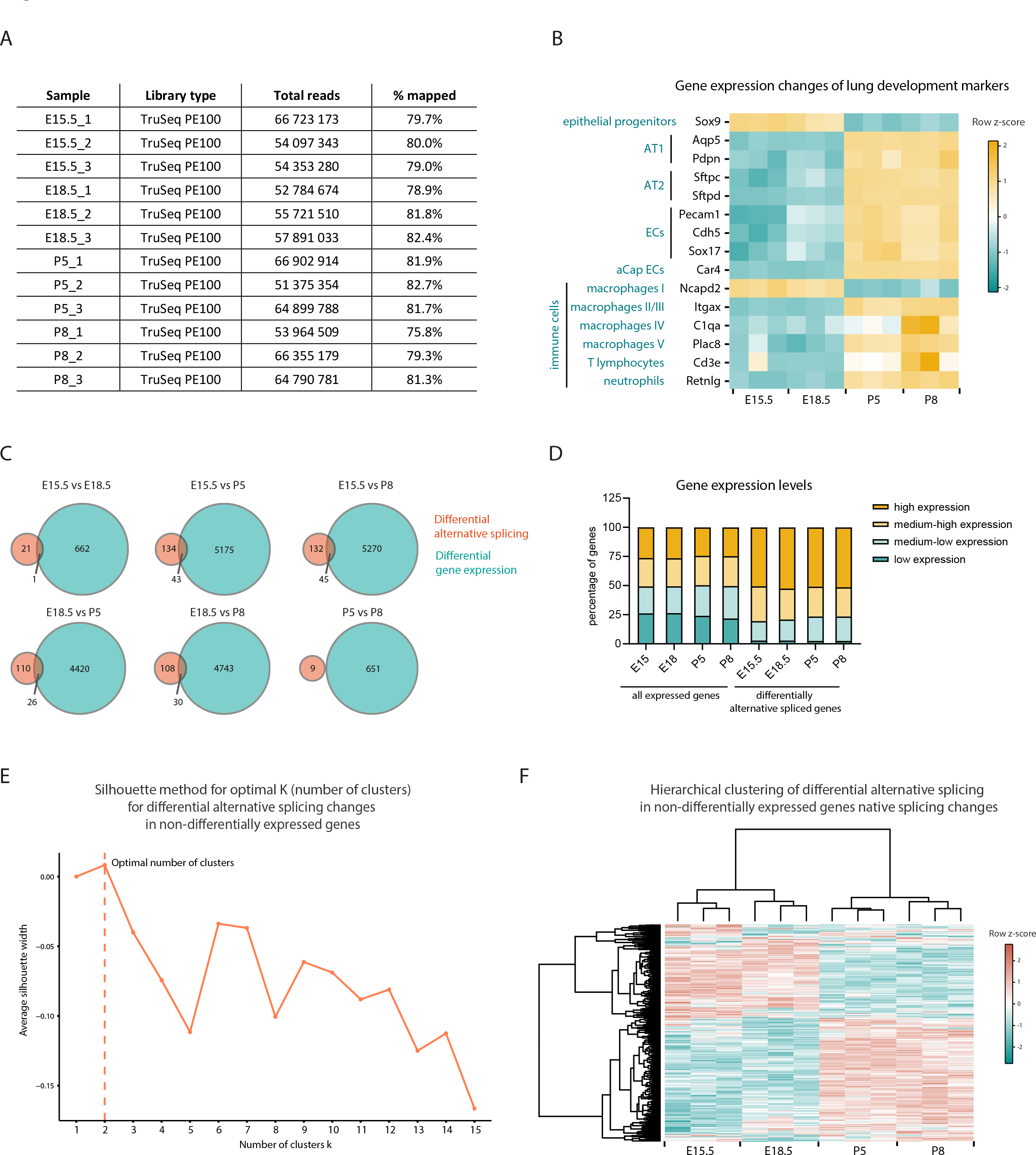
Analysis of RNAseq datasets from developing bulk mouse lungs. A. Total number of reads and percentage of alignment per sample. B. Heatmap representing gene expression changes of lung lineage-specific markers. Yellow and green represent increased and decreased gene expression, respectively, relative to the mean of each gene across the time course (row z-score calculated from log2(CPMs+1)). C. Venn diagrams representing the genes undergoing differential alternative splicing and/or differential gene expression in each pair-wise comparison between time-points associated. D. Distribution of gene expression levels of genes undergoing differential AS in at least one pair-wise comparison between time-points. The 16 152 expressed genes expressed in the mouse developing lungs were grouped in 4 bins of equal size according to their absolute level of expression (CPMs) (low expression, medium-low expression, medium-high expression, and high expression). Then, the genes undergoing differential AS in at least one pairwise comparison between time-points were distributed within these bins. E. Identification of the optimal number of clusters to use in K-means clustering of differentially AS events in at least one pair-wise comparison between time-points associated with non-differentially expressed genes in the same comparison using the average silhouette width method. The optimal number of clusters k is the one that maximizes the average silhouette, in this case K=2. F. Heatmap representing the hierarchical clustering of differentially AS events in at least one pair-wise comparison between time-points associated with non-differentially expressed genes in the same comparison. Cyan and coral represent decreased and increased PSI (percent spliced in), respectively, relative to the mean of each AS event across the time course (row z-score calculated from logit(PSI)).

**Figure S2.**
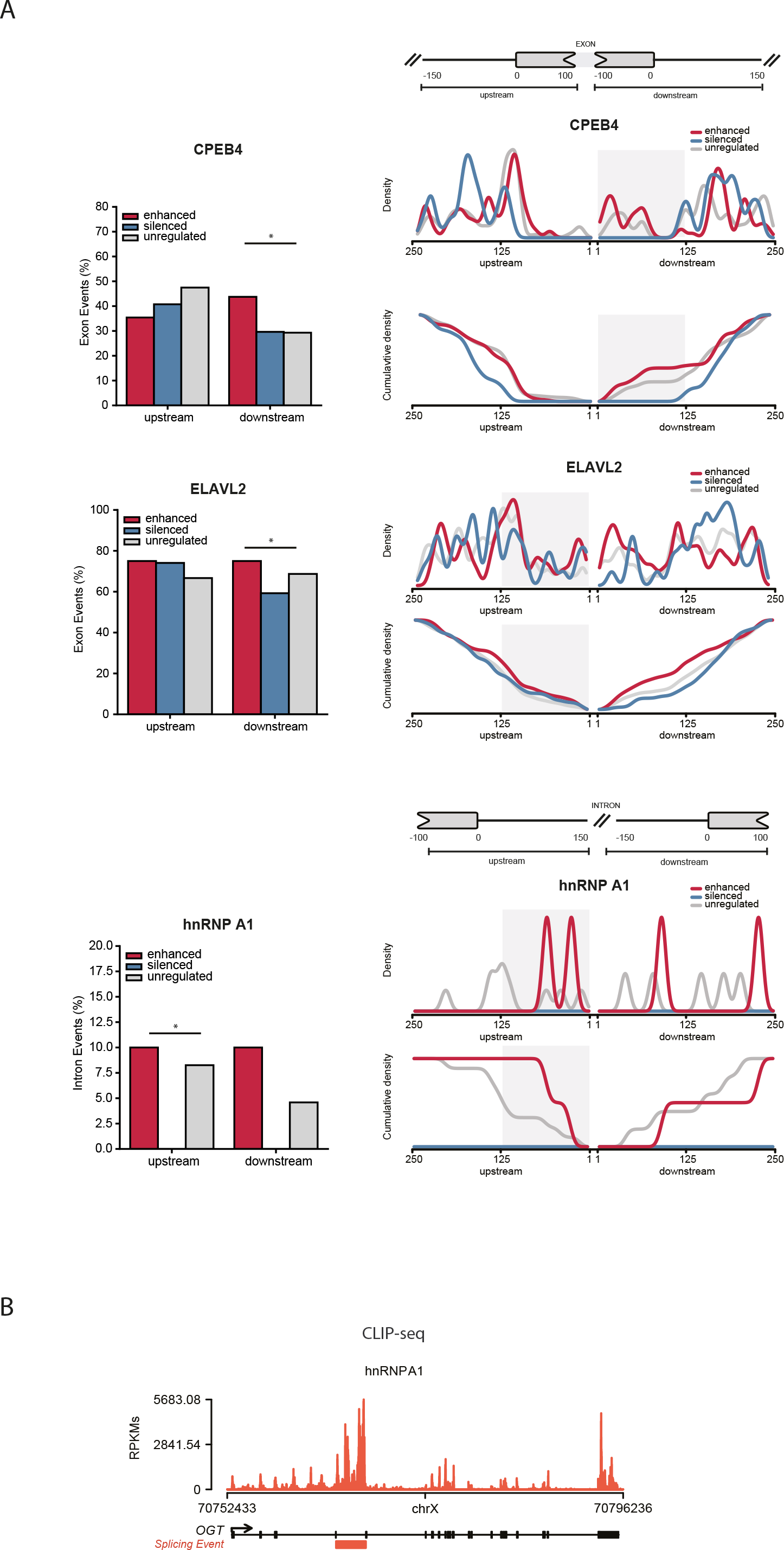
Candidate RBPs for the regulation of DAS between E18.5 and P5. A. Left: Bar plots show the relative number of events with at least one motif hit for each RBP normalized by the total number of respective events. Right: Density plots and cumulative density plots show the positional distribution of each RBP binding motif in regulated events against a background of unregulated events. These show where the motifs are more concentrated, and thus where the enrichment test was more significant (shaded area). Statistical significance was calculated using Matt test_regexp_enrich (p < 0.01). B. CLIP-seq signals at the genomic of the OGT represented as reads per kilobase million. The alternative splicing event is indicated in orange.

**Figure S3.**
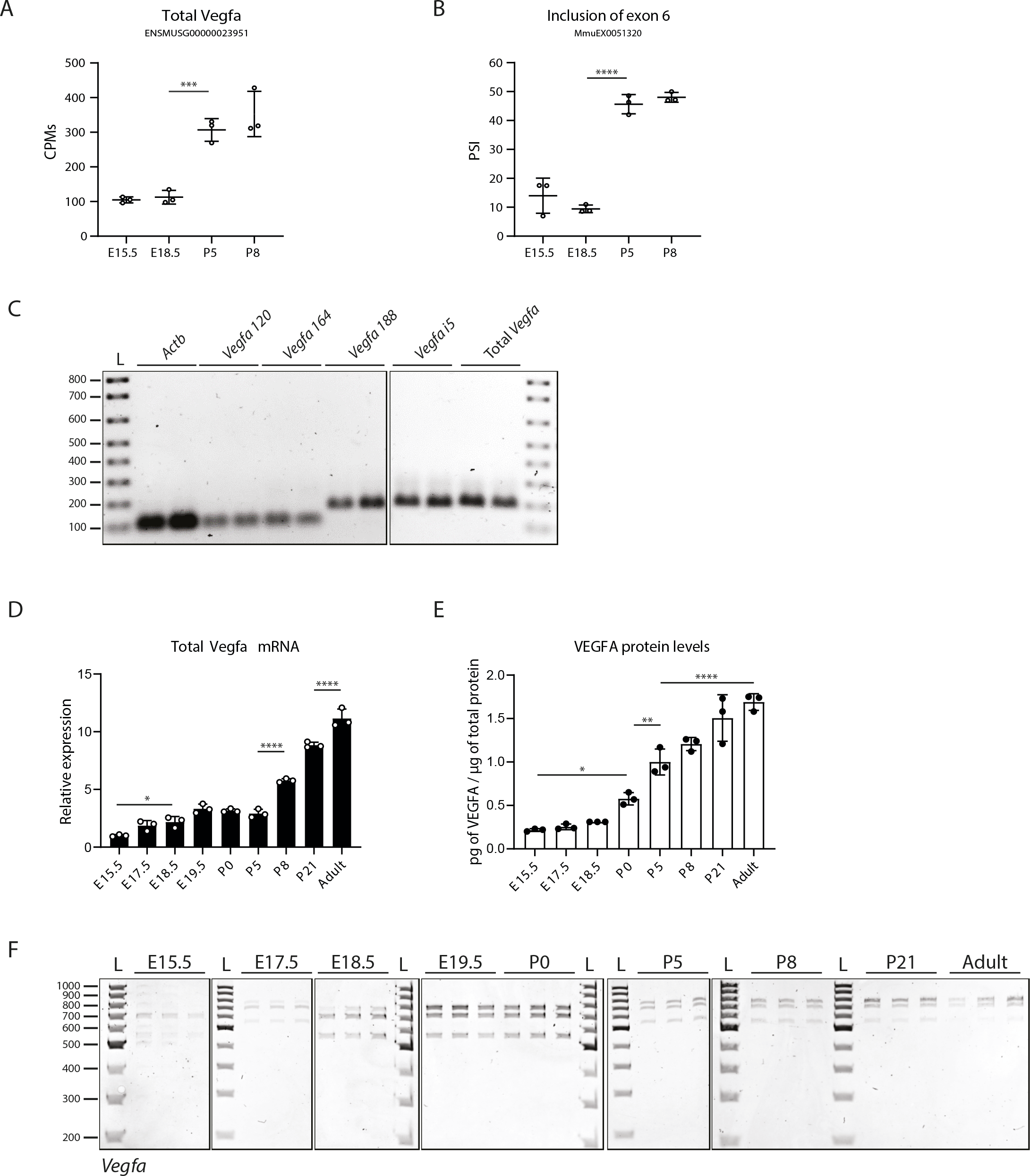
Analysis of *Vegfa* expression dynamics during lung development. A. Quantification of *Vegfa* gene expression levels from bulk lung RNAseq at different developmental time-points. CPM, Counts per million. p-value from unpaired t-test. B. Quantification of AS at different developmental time-points of a previously annotated alternatively spliced event of *Vegfa* associated with inclusion of exon 6 in Vast-TOOLS. The event is identified in Figure 3A. PSI, percent spliced-in. p-value from unpaired t-test. C. Analysis in agarose gels of the RT-qPCR amplification products obtained using the primer pairs indicated in Figure 4C. The presence of a single band of the correct size indicates that these primers pairs are specific. D. Expression changes of total *Vegfa* from bulk lungs at different developmental time-points analysed by RT-qPCR. N=3 for each time-point. E. VEGFA protein levels in bulk lungs quantified by ELISA. N=3 for each time-point. p-value from one-way ANOVA with Tukey correction for multiple testing. F. TBE-Urea PAGE gels used for the quantification of the normalized relative proportions between the expression levels of *Vegfa* isoforms on bulk lungs at different developmental time-points. N=3 for each time-point except for E19.5 (N=2).

**Figure S4.**
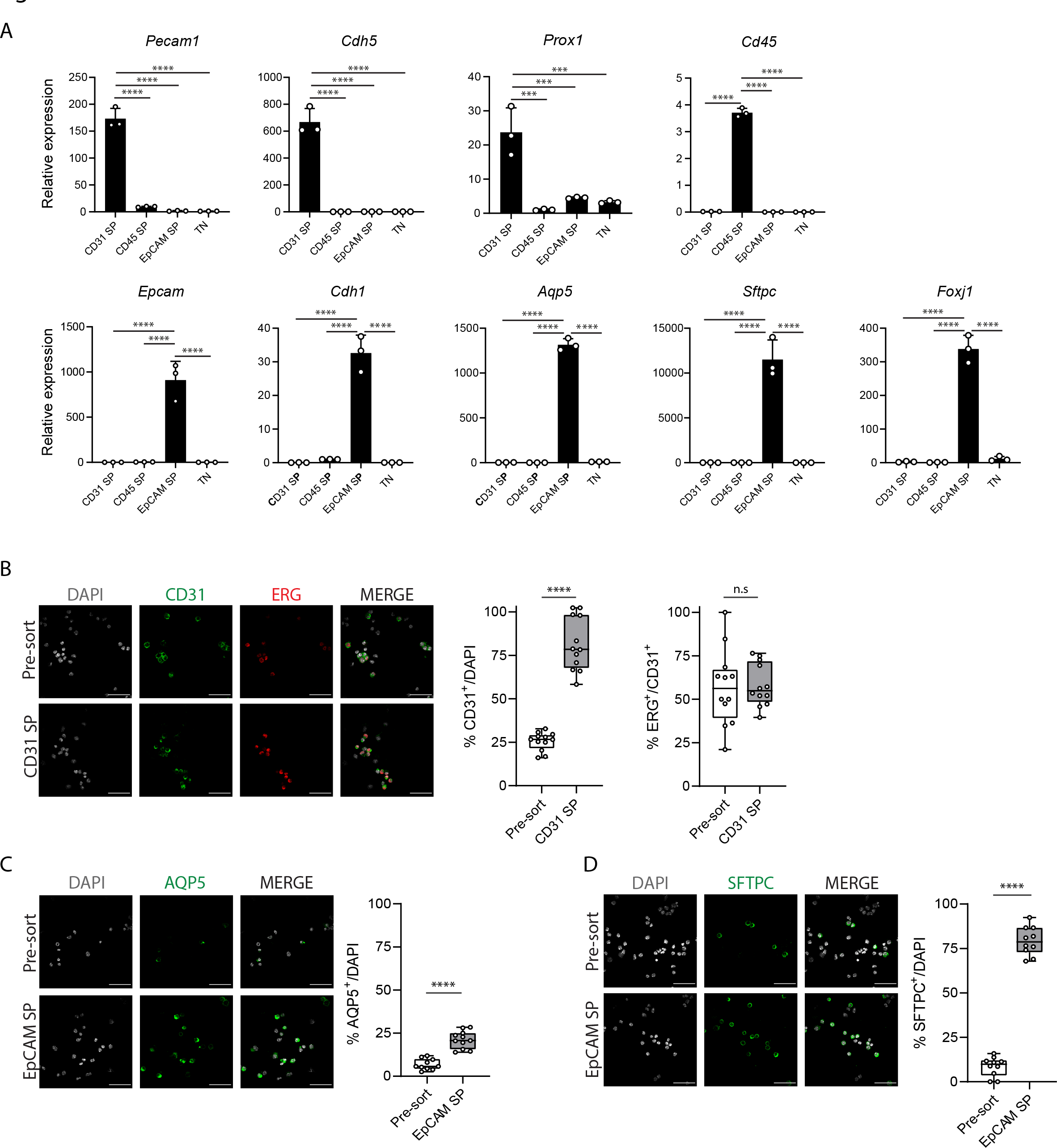
Isolation of distinct lung cell populations. A. Expression changes of cell type-specific markers from sorted cell populations at P5 by RT-qPCR. N=3 for each time-point. *Pecam1* (CD31) and *Cdh5* are markers for endothelial cells, *Prox1* is a marker for lymphatic endothelial cells, *Cd45* is a marker for immune cells, *Epcam* and *Cdh1* are markers for epithelial cells, *Aqp5* is a marker for epithelial AT1 cells, *Sftpc* is a marker for epithelial AT2 cells and *Foxj1* is a marker for epithelial ciliated cells. p-value from one-way ANOVA with Tukey correction for multiple testing. B. Left: Immunofluorescence of CD31 (green) and ERG (red) in pre-sort and sorted CD31 SP populations at P5. Cell nuclei are labeled with DAPI (grey). Scale bar, 22 um. Middle: Quantification of the percentage of CD31-positive cells in pre-sort and sorted CD31 SP populations at P5. Data from three independent experiments with at least 500 cells quantified per condition. Data are shown as mean ± SD. p-value from unpaired t-test. Right: Quantification of the percentage of ERG-positive cells from CD31-positive cells in pre-sort and sorted CD31 SP populations at P5. Data from three independent experiments with at least 500 cells quantified per condition. Data are shown as mean ± SD. p-value from unpaired t-test. C. Left: Immunofluorescence of AQP5 (green) in pre-sort and sorted EpCAM SP populations at P5. Cell nuclei are labeled with DAPI (grey). Scale bar, 22 um. Right: Quantification of the percentage of AQP5-positive cells in pre-sort and sorted EpCAM SP populations at P5. Data from three independent experiments with at least 500 cells quantified per condition. Data are shown as mean ± SD. p-value from unpaired t-test. D. Left: Immunofluorescence of SFTPC (green) in pre-sort and sorted EpCAM SP populations at P5. Cell nuclei are labeled with DAPI (grey). Scale bar, 22 um. Right: Quantification of the percentage of SFTPC-positive cells in pre-sort and sorted EpCAM SP populations at P5. Data from three independent experiments with at least 500 cells quantified per condition. Data are shown as mean ± SD. p-value from unpaired t-test.

**Figure S5.**
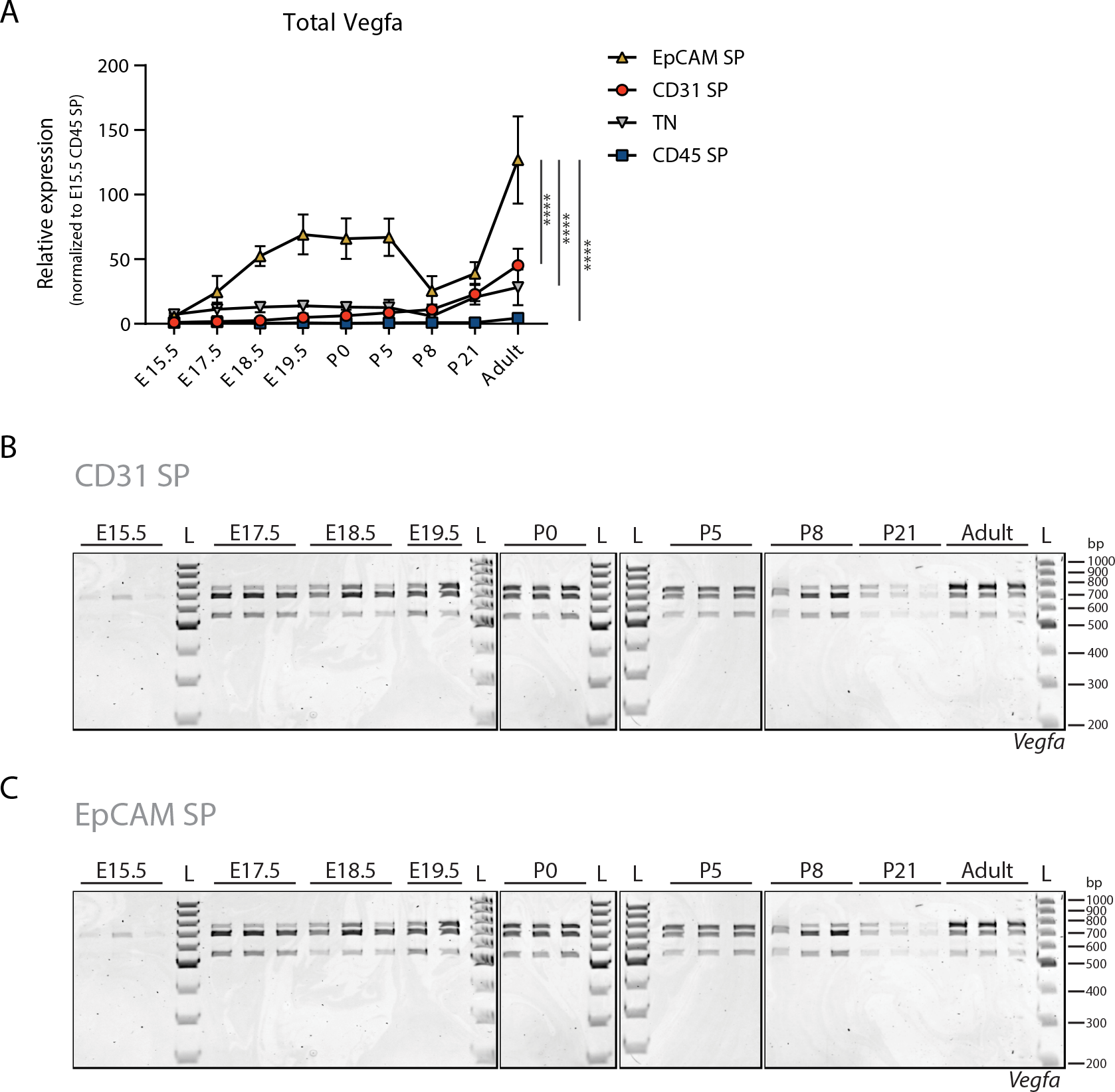
Analysis of *Vegfa* expression dynamics in distinct cell populations during lung development. A. Expression changes of Total *Vegfa* from sorted cell populations at different developmental time-points analysed by RT-qPCR. Expression values normalized to E15.5 for CD45 SP population. N=3 for each time-point. p-value from two-way ANOVA with Tukey correction for multiple testing. B. TBE-Urea PAGE gels used for the quantification of the normalized relative proportions between the expression levels of *Vegfa* isoforms on CD31 SP cell population at different developmental time-points. N=3 for each time-point. C. TBE-Urea PAGE gels used for the quantification of the normalized relative proportions between the expression levels of *Vegfa* isoforms on EpCAM SP cell population at different developmental time-points. N=3 for each time-point except E19.5 (N=2).

**Figure S6.**
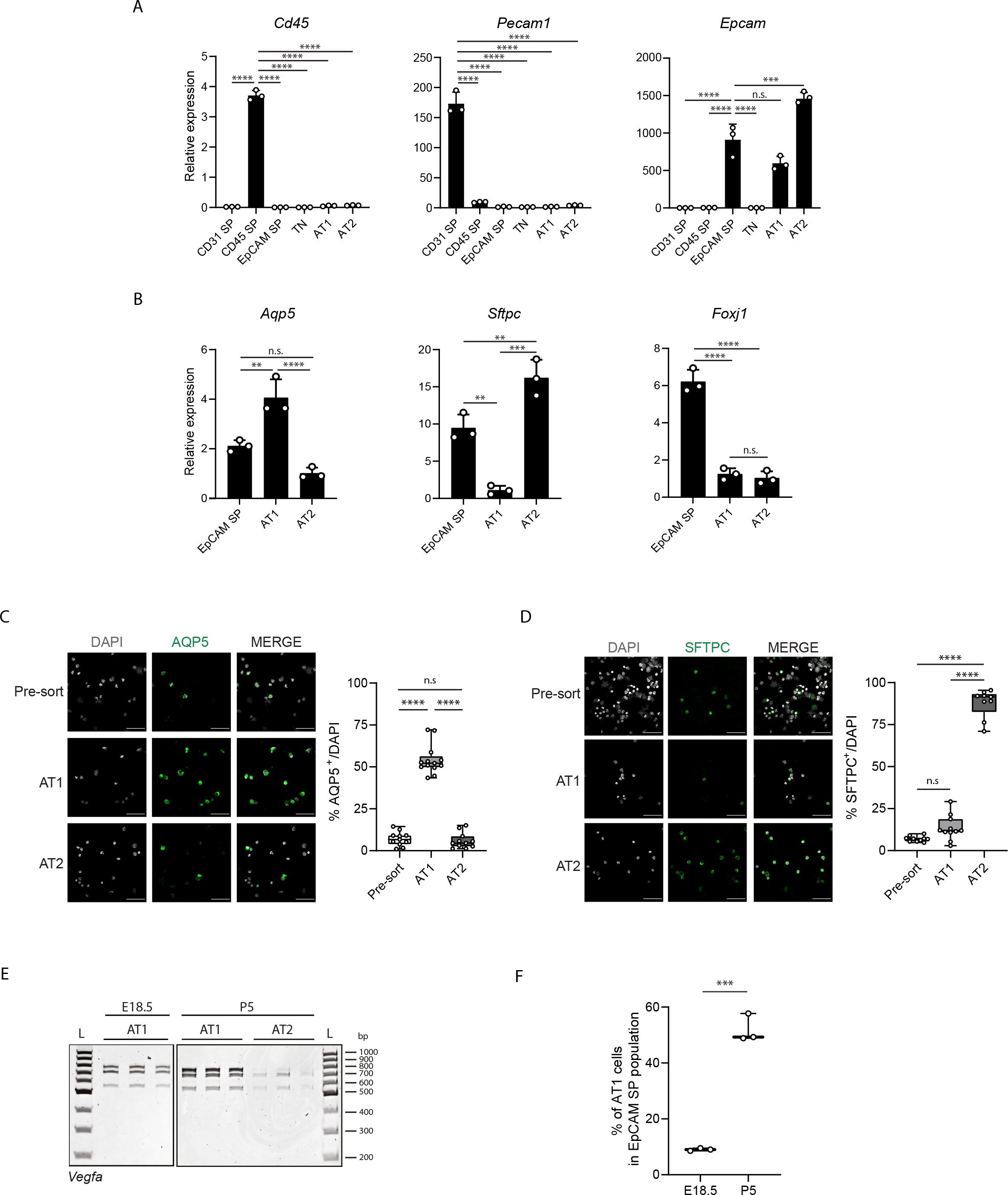
Isolation of distinct alveolar epithelial cell populations. A. Expression changes of cell type-specific markers from sorted cell populations at P5 by RT-qPCR. N=3 for each time-point. *Pecam1* (CD31) is a marker for endothelial cells, *Cd45* is a marker for immune cells, *Epcam* is a marker for epithelial cells. p-value from one-way ANOVA with Tukey correction for multiple testing. B. Expression changes of cell type-specific markers from sorted cell populations at P5 by RT-qPCR. N=3 for each time-point. *Aqp5* is a marker for epithelial AT1 cells, *Sftpc* is a marker for epithelial AT2 cells and *Foxj1* is a marker for epithelial ciliated cells. p-value from one-way ANOVA with Tukey correction for multiple testing. C. Left: Immunofluorescence of AQP5 (green) in EpCAM SP and sorted AT1 and AT2 populations at P5. Cell nuclei are labelled with DAPI (grey). Scale bar, 22 um. Right: Quantification of the percentage of AQP5-positive cells in EpCAM SP and sorted AT1 and AT2 populations at P5. Data from three independent experiments with at least 500 cells quantified per condition. Data are shown as mean ± SD. p-value from one-way ANOVA with Tukey correction for multiple testing. D. Left: Immunofluorescence of SFTPC (green) in EpCAM SP and sorted AT1 and AT2 populations at P5. Cell nuclei are labelled with DAPI (grey). Scale bar, 22 um. Right: Quantification of the percentage of SFTPC-positive cells in EpCAM SP and sorted AT1 and AT2 populations at P5. Data from three independent experiments with at least 500 cells quantified per condition. Data are shown as mean ± SD. p-value from one-way ANOVA with Tukey correction for multiple testing. E. TBE-Urea PAGE gels used for the quantification of the normalized relative proportions between the expression levels of *Vegfa* isoforms at E18.5 on AT1 and for AT1 and AT2 cell populations at P5. N=3 for each time-point. F. Quantification of the FACS analysis of the fractions of EpCAM^low^ MHC II^-^ cells (AT1) within the EpCAM-positive population extracted from mouse lungs at E18.5 and P5. N=3 for each time-point. p-value from unpaired t-test.

## Supplemental tables

Table S1. DAS events in at least one pairwise comparison between time-points, as calculated by Vast diff (|ΔPSI|>10%, confidence interval 95%).

Table S2. Differentially expressed genes in at least one pairwise comparison between time- points, as calculated by EdgeR (|log2FC|>1, FDR<0.05).

Table S3. DAS events in at least one pairwise comparison between time-points associated with non-differentially expressed genes in the same comparison.

Table S4. DAS events in at least one pairwise comparison between time-points associated with differentially expressed genes in the same comparison.

Table S5. DAS events occurring in non-differentially expressed genes included in each kinetic cluster from K-means clustering and associated PSIs for each time-point.

Table S6. Enriched KEGG pathway terms for DAS events occurring in non-differentially expressed genes

Table S7. RNA-binding motifs enriched in DAS events occurring in non-differentially expressed genes between E18.5 and P5

Table S8. RBPs identified to undergo differential gene expression between E18.5 and P5.

Table S9. DAS events between E18.5 and P5 that do not undergo differential gene expression in this same time interval (DAS_nonDEG) with enriched motifs for each differentially expressed RBP.

Table S10. DAS events between E18.5 and P5 that undergo differential gene expression in this same time interval (DAS_DEG) with enriched motifs for each differentially expressed RBP.

Table S11. RBPs motifs enriched along Vegfa intron 5 sequence

## Materials and Methods

### Key resources table

| Reagent or Resource | Source | Identifier |
| --- | --- | --- |
| **Antibodies** |  |  |
| Rat anti-mouse APC CD31 | BD Pharmingen | 551262 |
| Mouse anti-mouse FITC CD45 | Covance | 109806 |
| Rat anti-mouse PE CD326 (Ep-CAM) | Covance | 118206 |
| Rat anti-mouse PerCP/Cy5.5 MHC II | Biolegend | 107626 |
| Pro-SFPTC, Rabbit | SevenHills BioReagents | WRAB-9337 |
| CD31 (PECAM1), Goat | R&D | AF3628 |
| Aquaporin 5, Rabbit | Merck | 178615 |
| ERG, Rabbit | Abcam | ab92513 |
| Donkey anti-Rabbit Alexa 568 | Thermo Fisher Scientific | A10042 |
| Donkey anti-goat Alexa 647 | Thermo Fisher Scientific | A21447 |
| **Chemicals, Peptides and Recombinant Proteins** |  |  |
| PBS | Sigma-Aldrich | P4417 |
| Dulbecco's Modified Eagle Medium (DMEM) | Gibco | 11320-074 |
| Albumin Bovine Fraction V (BSA) | Nzytech | MB04602 |
| RNase-OFF | Enzifarma | 9037 |
| Liberase™ Research Grade | Roche | 5401119001 |
| DNase I, grade II, from bovine pancreas | Roche | 10104159001 |
| Ethylenediaminetetra-acetic acid (EDTA) | VWR | VWRC20301.290 |
| RBC lysis buffer 10x | Biolegend | 420301 |
| RNase-free water | Sigma-Aldrich | 3098 |
| Trypan Blue | Sigma-Aldrich | T8154 |
| Live/Dead Fixable Near-IR Dead cell Stain kit | Invitrogen | LTI L10119 |
| Hepes | Sigma-Aldrich | H3375-250G |
| TRIzol™ Reagent | Alfagene/Ambion | 15596026 |
| Chloroform | Merck Millipore | 1.02445.1000 |
| Glycogen | Sigma | G1767-1VL |
| 3M Sodium Acetate | VWR | 27653,26 |
| Ethanol Absolute | VWR | 20821,33 |
| Phenol-Chloroform-Isoamyl alcohol (25:24:1) | Sigma | P2069 |
| Isopropanol | VWR | 1.09634.1000 |
| Power SYBR® Green PCR Master Mix | Thermo Fisher sci | 4368706 |
| Tris base | Nzytech | MB01601 |
| Agarose | Nzytech | MB14402 |
| GreenSafe Premium | Nzytech | MB13201 |
| DNA Gel Loading Dye (6X) | Thermo | R0611 |
| GeneRuler 1Kb Plus DNA ladder | Thermo Scientific | SM1333 |
| GeneRuler 100 bp DNA Ladder | Thermo | SM0241 |
| Urea | Sigma-Aldrich | U5378 |
| Ammonium Persulfate | Sigma | A3678-25G |
| Ficoll 400 | Enzymatic | BP525-25 |
| Sodium chloride | VWR | 27810,295 |
| Triton X-100 | Sigma-Aldrich | T8787 |
| Dithiothreitol (DTT) | Enzo Life Sciences | ALX 280 001 G010 |
| Halt™ Protease and Phosphatase Inhibitor Cocktail | Thermo | 78446 |
| Acrylamide/bis-Acrylamide (29:1 solution) | Nzytech | MB04501 |
| Hydrochloric acid | VWR | 20252,335 |
| N,N,N',N'-TETRAMETHYLETHYLENEDIAMINE | Sigma | T9281-25ML |
| Bromophenol Blue | Merck | 108122 |
| RNeasy Plus Mini kit | Qiagen | 50974136 |
| RNeasy Plus Micro kit | Qiagen | 50974034 |
| DNase I recombinant, RNase-free | Roche | 4716728001 |
| High Capacity RNA-to-cDNA Kit | Applied Biosystems | LTAB 4387406 |
| Q5 Hot Start High-Fidelity DNA Polymerase | BioLabs | M0493S |
| Wizard® SV Gel and PCR Clean-Up System | Promega | A9281 |
| Pierce™ BCA Protein Assay Kit | Thermo Scientific | 23227 |
| Mouse VEGFA Quantikine® Elisa kit | R&D | MMV00 |
| **Experimental Models: Organisms** |  |  |
| C57Bl/6J mice (*Mus musculus* Linnaeus) | Charles River Laboratoires | N/A |
| **Oligonucleotides** |  |  |
| Primers used in PCR are listed in the methods section | Sigma-Aldrich | This manuscript |
| **Bioinformatics tools and softwares** |  |  |
| Kallisto |  | https://github.com/pachterlab/kallisto |
| VastDB |  | https://vastdb.crg.eu/wiki/Main_Page |
| VAST-TOOLS |  | https://github.com/vastgroup/vast-tools |
| Matt |  | http://matt.crg.eu/ |
| DAVID |  | https://david.ncifcrf.gov/tools.jsp |
| QuantStudio™ Real-time PCR Software | Applied Biosystems | 4486400 |
| Prism 8 Software | GraphPad | https://www.graphpad.com/scientific-software/prism/ |
| Image Lab™ Software | BioRad | 1709690 |
| SnapGene® Software | GSL Biotech | https://www.snapgene.com |
| FLOW JO Software | BD Life Sciences | https://www.flowjo.com/ |
| ZEN software (black edition) | Carl Zeiss | N/A |
| ImageJ Software | NIH | https://imagej.nih.gov/ |
| Microsoft Excel 2013 | Office | N/A |
| **Others** |  |  |
| **Sanger Sequencing** | Eurofins \|genomics /GATC services | N/A |

### Mice and sample collection

Wild-type C57Bl/6 mice (*Mus musculus Linnaeus*) were used in this study and were maintained under normal husbandry conditions at the Instituto de Medicina Molecular (iMM) and under national regulations. Animal procedures were performed under the DGAV project license 0421/000/000/2016. Both female and male mice were used for sample collection.

Timed matings were performed and mouse lungs were collected at different time-points (E17.5, E18, E18.5, E19, E19+2h, E19.5, P0, P0+2h, P2 and P8) for different procedures. Biological replicates of each time-point were collected from the same litter.

### RNA extraction from bulk lungs

Immediately after isolation, lungs were transferred to an Eppendorf, snap frozen in liquid nitrogen and stored at -80°C. The lungs were thawed and mechanically disrupted using a pestle. To extract and purify RNA, we used the RNeasy Plus Mini Kit (Qiagen), following manufacturers’ instructions. For further dissociation of the tissue, after RLT plus lysis buffer addition, we used the pestle and a 20G needle in a 3 mL syringe (passing the tissue roughly 10 times). RNA concentration was measured using Thermo Scientific™ NanoDrop 2000. A fraction of the purified RNA was used to produce cDNA and the remaining volume of RNA was stored at -80°C.

### Lung tissue dissociation and FACS

For cell sorting, the lungs were collected and transferred to a 50 mL falcon with 1 mL of cold (+4°C) Dulbecco’s Modified Eagle Medium (DMEM) supplemented with 1% Bovine Serum Albumin (BSA). Disruption of the tissue was performed on ice using sterile scissors and a sterile scalpel blade until no clear tissue pieces were visible. The dissociated lung tissue was incubated with 2 mL of Liberase (200 µg/mL) and DNase I (10 µg/mL) in DMEM + 1% of BSA in an Eppendorf tube and incubated in a rotator for 40 min at 37°C. During this enzymatic digestion period, the tissue was further disrupted mechanically by passing 4 times through a 20G needle attached to a 2 mL syringe. We added 2 mL of cold FACS buffer (EDTA 1 mM, BSA 0.5%, dPBS 1x) and aspirated the suspension into a 10 mL syringe with a 20G needle attached. Cell suspension was forced to pass through a 70 µm cell strainer. We centrifuged the samples at 200 x g for 5 min at 4°C. The supernatant was discarded. To lyse red blood cells (RBC), each pelleted lung was re-suspended in 1 mL of RBC lysis buffer 1x and incubated for 5 min at RT. Cell suspension was forced to pass through a 40 µm cell strainer. The number of cells was counted and the cell suspension was centrifuged at 200 x g for 5 min at 4°C. The pelleted cells were re-suspended in FACS buffer to a density of 16x10^6^ cells/mL. Incubation with antibodies and LiveDead Fixable Viability dye (L/D) was performed for 20 min in the dark at +4°C in the following dilutions: Live/Dead APC-Cy7 (1:2000); Rat anti-mouse APC CD31 (1:100); Mouse anti-mouse FITC CD45 (1:200); Rat anti-mouse PE CD326 (Ep-CAM) (1:100); and Rat anti- mouse PerCP/Cy5.5 MHC II (1:500). Single color and unstained controls were used to set up the gatings.

After antibody incubation, cells were centrifuged at 300 x g for 5 min at 4°C and washed by re-suspending in FACS buffer to a concentration of 1x10^6^ cells/200 µL and centrifugation at 300 x g for 5 min at 4°C. Finally, pelleted cells were re-suspended in FACS buffer to a final concentration of 1x10^6^ cells/250 µL for the single colors, and unstained fraction, and 8x10^6^ cells/mL for the fraction to be sorted. FACS was performed in BD FACSAria III cell sorter with a nozzle of 100 µm and a pressure of 20 PSI. Dead cells were excluded using LiveDead Fixable Viability dye. We collected different cell populations, in RNase-free Eppendorf tubes filled with 500 µL of collection buffer (Hepes 25 mM, BSA 2.5% in DMEM) and centrifuged at 2400 x g for 5 min at 4°C. The supernatant was discarded and the pellet was re-suspended in 1 mL of Trizol Reagent by pipetting up and down several times vortexed for 1 min. After homogenization, cells were incubated at RT for 5 min to allow the complete dissociation of nucleoprotein complexes. Cell lysates were stored at -80°C until RNA extraction of sorted cells was performed.

### RNA extraction from sorted cells

The samples in Trizol were thawed on ice and incubated at RT for 5 min. 200 µL of chloroform was added to each Eppendorf. The tubes were shaken by hand for 30 sec and incubated at RT for 5 min. After incubation, the tubes were centrifuged at 12 000 x g for 15 min at 4°C. The upper aqueous phase was carefully transferred to a new RNase-free Eppendorf tube. To precipitate and wash RNA, 1.5 µL of glycogen and 50 µL of 3M sodium acetate were added to each tube. Tubes were briefly vortexed and 500 µL of isopropanol was added. The eppendorfs were vortexed and incubated at RT for 15 min and centrifuged at 12 000 x g for 8 min at 4°C. The supernatants were discarded and the pellets were washed with 1 mL of 75% ethanol. The tubes were centrifuged at 12 000 x g for 5 min at 4°C and the supernatant was carefully discarded after centrifugation. The pellets were allowed to dry at RT and were resuspended in 20 µL of RNase-free water. After suspension, all samples were kept on ice and RNA was quantified using Thermo Scientific™ NanoDrop 2000. Samples were treated with recombinant DNase I (RNase-free). For each reaction 5 µL of DNase I buffer, 1 µL of DNase I and 24 µL of RNAse-free water were added to 20 µL of RNA (up to 10 µg) from each sample making a final volume of 50 µL per reaction and incubated for 20 min at 30°C. To inactivate DNase I and purify the RNA, we added 1 volume of phenol-chloroform-isoamyl alcohol mixture (25:24:1). The tubes were vortexed and centrifuged at 12 000 x g for 10 min at 4°C. The upper aqueous phase was transferred to a new RNase-free Eppendorf tube. 1 volume of chloroform was added to each sample, the samples were then vortexed and centrifuged at 12 000 x g for 10 min at 4°C. The upper aqueous phase was transferred to a new RNase-free Eppendorf. To precipitate RNA, we added 1.5 µL of glycogen and 50 µL of 3M sodium acetate to each tube and vortexed them. 500 µL of isopropanol was added, the tubes were vortexed and incubated at RT for 15 min. After incubation, the tubes were centrifuged at 12 000 x g for 20 min at 4°C to precipitate RNA and the supernatant was discarded. To wash the pellets, we added 1 mL of 75% ethanol and centrifuged the tubes at 12 000 x g for 5 min at 4°C. We discarded the supernatant carefully and repeated this step once to further wash the RNA. Pellets were dried at RT and re-suspended it in 15 µL of RNase-free water. Each sample was kept on ice and quantified using Thermo Scientific™ NanoDrop 2000. A fraction of the purified RNA was used to produce cDNA and the remaining volume of RNA was stored at -80°C.

### Production of cDNA

Production of cDNA from RNA extracted from both bulk lungs and sorted cells was performed using the High-Capacity RNA-to-cDNA™ Kit (Applied Biosystems™), following the manufacturers’ protocol. The cDNA was then stored at -20°C and used for end-point PCR and RT-qPCR reactions.

### End-point Polymerase Chain Reaction (PCR)

Specific target regions were amplified through standard end-point PCR. Primers used to amplify the distinct *Vegfa* isoforms through end-point PCR are presented in the table below. PCR was performed using Q5 Hot Start High-Fidelity DNA Polymerase. To perform each reaction we added: 10 µL of 5X Q5 Reaction Buffer, 1 µL of dNTPs (10 mM), 2.5 µL of Forward Primer (10 µM), 2.5 µL of Reverse Primer (10 µM), 0.5 µL of Q5 Hot Start HighFidelity DNA Polymerase, cDNA as template and Nuclease-Free water up to 50 µL. PCR was run on T100™ Thermal Cycler (Bio-Rad) using the following PCR program: an initial denaturation step of 98°C for 30 secs, 26 cycles of 10 sec at 98°C, 30 sec at 65°C and 2 min at 72°C, followed by a final extension of 2 min at 72°C. PCR products were analysed by performing electrophoresis in a 2% agarose gel or in a TBE-UREA-Polyacrylamide gel as described below.

### Primers used in standard end-point PCR

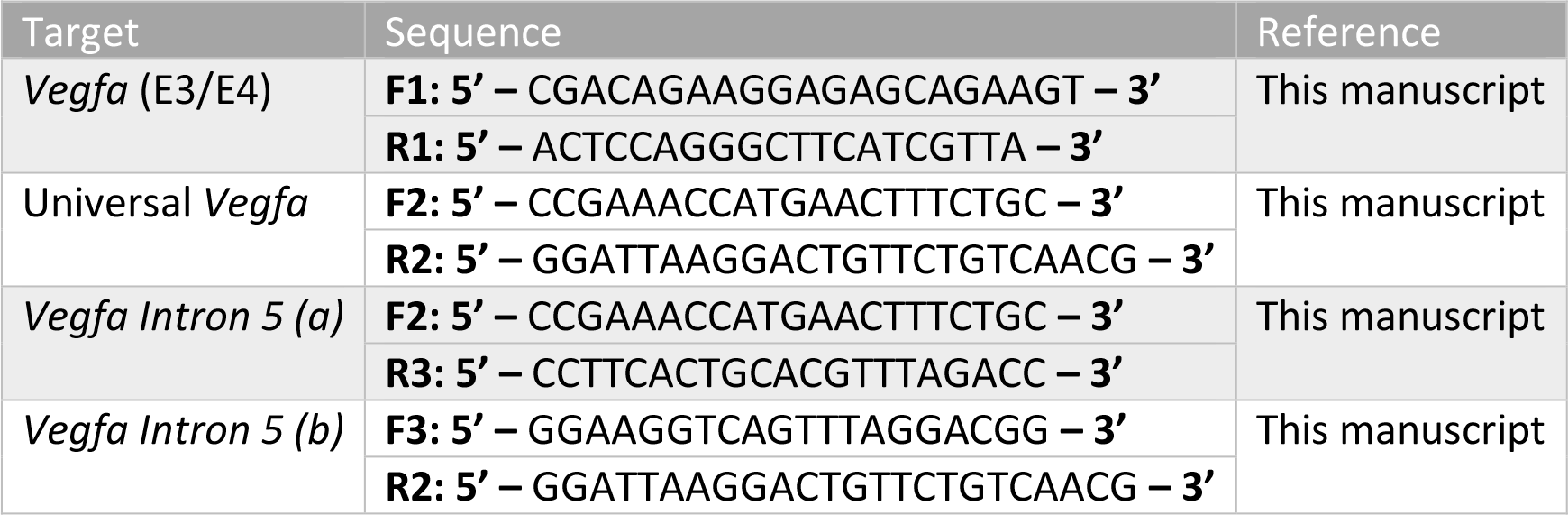

### PCR product purification and Sanger sequencing

To sequence the *Vegfa* isoform comprising intron 5, several overlapping sequences were obtained in a standard end-point PCR reaction. From the total volume of PCR product, we saved a fraction (10 µL) to run on a 2% agarose gel to confirm if the amplification of each fragment was successful. The remaining volume in the tube (40 µL) was purified using the Wizard® SV Gel and PCR Clean-Up System (Promega) following the manufacturers’ protocol. DNA Sanger Sequencing was performed by GATC services (Eurofins|genomics). DNA sequences obtained from the sequencing results of fragments amplified from *Vegfa intron 5* (FASTA file) were imported into the SnapGene® software (GSL Biotech).

### Gene expression analysis using real-time quantitative PCR (RT-qPCR)

Quantitative gene expression analysis was performed by RT-qPCR using intron-spanning primer pairs. Two housekeeping genes, *Actb* and *Gusb*, were analysed as housekeeping genes. All primers used for RT-qPCR are indicated in the table below. A standard curve for each primer pair was obtained in every RT-qPCR run alongside with the samples to be analysed. To obtain the standard curve, we mixed cDNA from all time-points collected and prepared different dilutions (1:10, 1:100, 1:500, 1:1000).

### Primers used in RT-qPCR

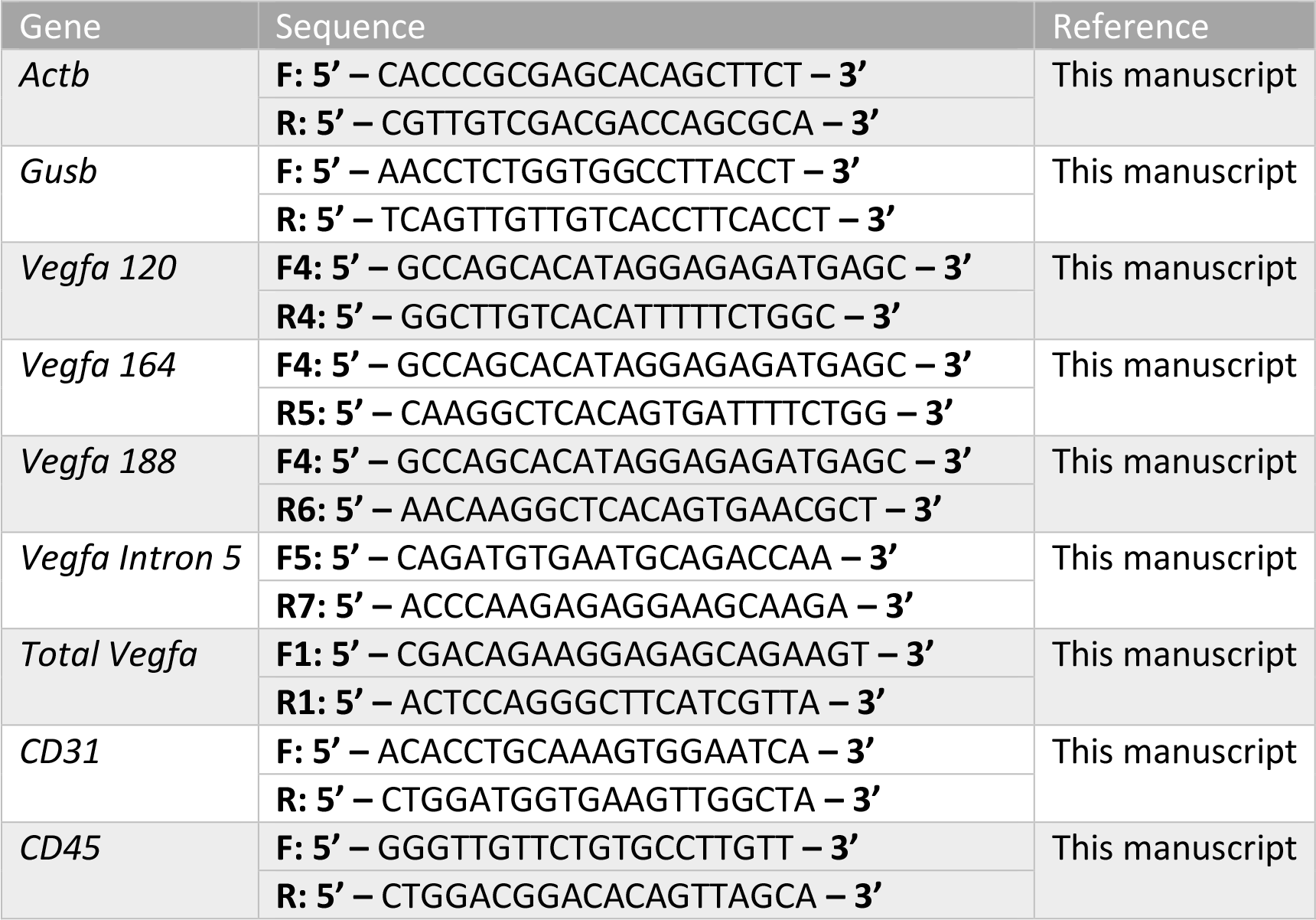
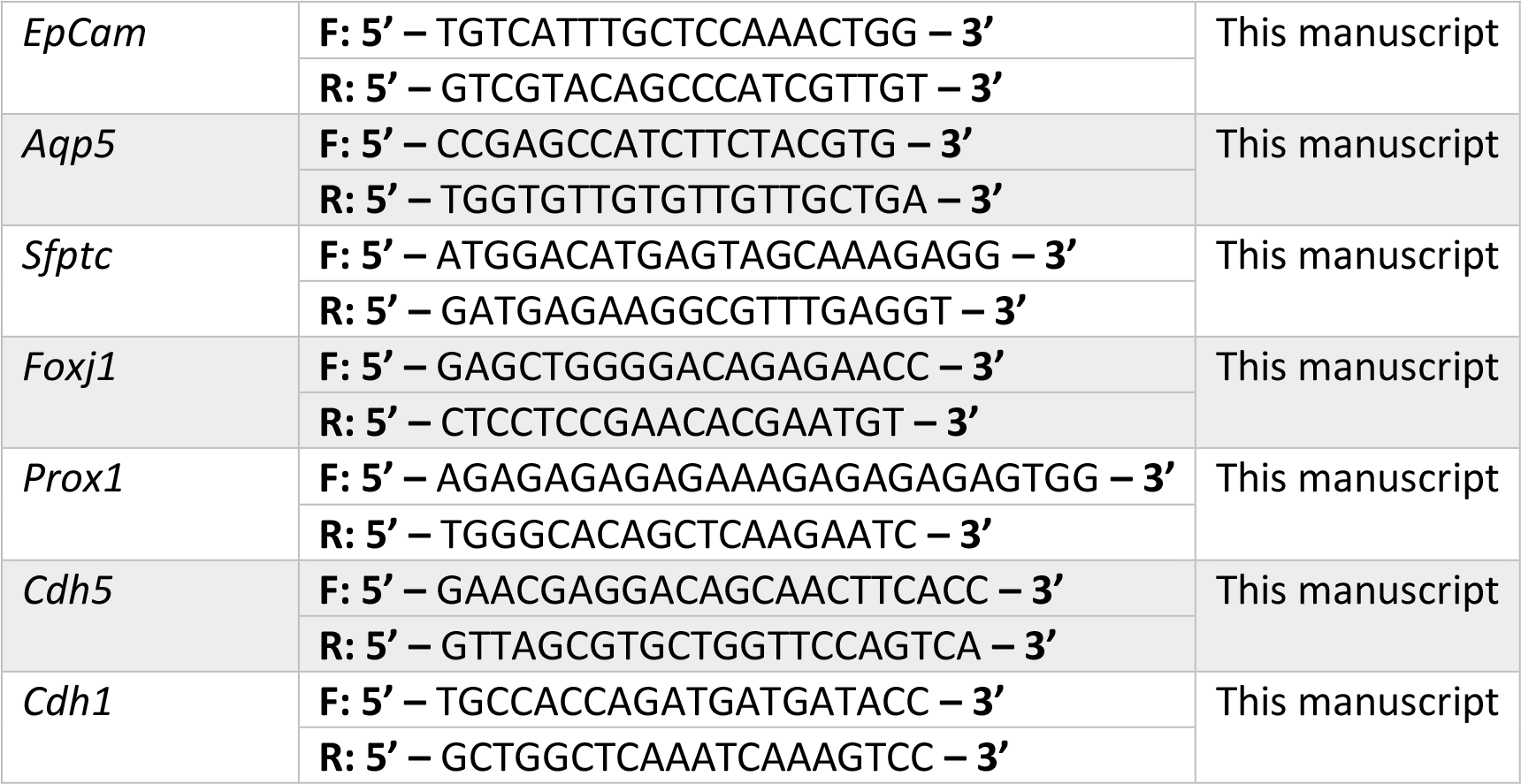

Additionally, in each plate we used a calibrator sample. The calibrator sample, which was prepared once and used in every plate, was generated by mixing equal quantities of each cDNA sample (all time-points) from bulk lungs (1:200). For each individual reaction we added: 7 µL of Power SYBR® Green PCR Master Mix, 0.3 µL of previously diluted primer pairs (final concentration of 100 nM), 2 µL of diluted cDNA and 4.85 µL of RNase-free water resulting in a final volume of 14 µL per well. We used Applied Biosystems VIIA 7 Real-Time PCR system, the conditions for the reaction were: 1x 50°C for 2 min, 95°C for 10 min; 45x 95°C for 15 sec,60°C for 1 min, 1x 95°C for 15 sec, 60°C for 1 min; and 1x 95°C for 15 sec. The software used to analyze each RT-qPCR experiment was QuantStudio™ Real-time PCR Software (Applied Biosystems). Standard curves, Melting curve and amplification plots were all generated in this software. Pfaffl method ^58^ was used to quantify gene expression. To avoid inter-plate variations when we needed to compare between different plates, we used an adaptation of the method used in qbase+ software (Biogazelle) that allows relative quantification of gene expression through a modified method based on ΔΔCt and through the normalization to a calibrator sample (Calibrated Normalized Relative Quantity, CNRQ). All graphics presented were elaborated with Graph Pad Prism 8 software.

### TBE-Urea-Polyacrylamide gel electrophoresis (PAGE)

6% of TBE-Urea-Polyacrylamide gels were prepared by using: 1.5 mL of 10x TBE Buffer, 2.25 mL of 40% polyacrylamide/bisacrylamide reagent, 7.2 g of Urea and ddH2O up to 15 mL. After completely dissolving the urea, we added 15 µL of TEMED and 150 µL of fresh 10% (w/v) APS. The mixture was poured into a 15mm-thick gel support and a 15-well comb was inserted. After polymerization, we mounted the gels in an electrophoresis apparatus filled with 1x TBE buffer. Before loading the samples, urea traces and gel pieces were washed from the wells with 1x TBE Buffer and the gel was pre-ran for 30 min at 25V. We loaded 30 µL of sample in each well - 15 µL of PCR product diluted in 15 µL of homemade 2x TBE-Urea sample buffer (10% 10x TBE running Buffer; 6% Ficoll Type 400; 1% bromophenol blue; 7M Urea). The gel was run at 25V until the samples moved past the loading space into the gel, after which the voltage to 100V was increased. After running, the gel was incubated in 1x TBE for 10 min and after was stained with Green Safe reagent dye (5 µL /100 mL of TBE 1x) for 30 min and visualized on ChemiDoc XRS+ system (BioRad).

### Cytospin and immunofluorescence

Single cell suspension from lungs at P5 before sorting (pre-sort sample) and after sorting were used in cytospin ^40^. 80 000 -100 000 cells/ 200 uL FACS buffer were used per slide. Samples were centrifuged in Shandon Cytospin 2 for 5 min at 500 g. The resultant slide was fixed with 4% PFA for 10 min at RT, washed twice in PBS and stored in PBS 0.01% Azide at +4°C until further use.

Immunostaining was performed in cytospin slides (all steps at RT). Slides were blocked in 3%BSA 0.1% triton X-100 in PBS (PBST-BSA) for 30 min. Primary antibody incubation was performed in PBST-BSA for 2 hours. Slides were washed three times in PBS 0.1% Triton X-100 (PBST). Secondary antibody incubation was performed in PBST-BSA for 1h. Slides were washed three times in PBST. Nuclei were stained with DAPI form 5 min. Slides were washed once with PBS and mounted in Mowiol/Dabco mixture. Imaging was performed with a 40x EC Plan-Neofluar DIC objective. Image analysis was performed with Fiji software.

The antibodies used in immunofluorescence were: Rabbit Pro-SFPTC (SevenHills BioReagents, 1:1000), Goat CD31 (PECAM1) (R&D, 1:400), Rabbit Aquaporin 5 (Merck, 1:200), Rabbit ERG (Abcam, 1:200), Donkey anti-Rabbit Alexa 568 (Thermo Fisher Scientific, 1:500), and Donkey anti-goat Alexa 647 (Thermo Fisher Scientific, 1:500)

### ELISA

After collection, lungs were snap-frozen in liquid nitrogen. We added cold lysis buffer (150 mM NaCl; 1 mM EDTA; 50 mM Tris-HCl pH=7.4; 1% Triton X-100 diluted in ddH2O and supplemented with 1 mM DTT and proteinase and phosphatase inhibitors (Thermo Scientific, 1861282) to the tube with the frozen lungs. Volume of lysis buffer was adjusted according to lung size (500 µL/150 mg). To mechanically disrupt the tissue, we used a pestle and pipetted up and down. We incubated the mixture on ice for 15 min and centrifuged at maximum speed for 15 min at +4ᵒC. We transferred the supernatant to a new ice-cold tube. To quantify protein concentration, we used the Pierce BCA Protein assay kit following the manufacturers’ protocol. For analysis of VEGFA protein levels, we used the Mouse VEGF Quantikine ELISA Kit (R&D, MMV00), following the manufacturers’ protocol.

### RNAseq and bioinformatics analysis

#### RNA isolation and sequencing library preparation

We performed RNAseq using mouse lung at 4 developmental stages (E15.5, E18.5, P5 and P8) in triplicates. After RNA extraction, RNA integrity was evaluated in Fragment Analyzer and all RNA samples revealed to have RNA quality number (RQN) >9.8. Library preparation was performed by using Truseq RNA Library protocols. Samples were barcoded, pooled and redistributed into 3 lanes. RNA sequencing was performed using HiSeq 4000 sequencing platform. In total, we obtained 897 million paired-end (PE) 101 nt reads (59.8 million per sample on average) **(Figure S1A)**. The RNAseq data has been deposited into the NCBI Gene Expression Omnibus (https://www.ncbi.nlm.nih.gov/geo/) under the accession number GSE175403.

#### RNAseq data alignment and differential gene expression

The ∼56.8 M raw reads generated per sample were uniquely mapped to the mm10 assembly and annotated to the Gencode vM14 transcriptome using TopHat2 (version 2.1.1) ^59^, with Bowtie2 (version 2.3.4) ^60^.

For gene expression analysis, gene expression levels were quantified with HTSeq-count (version 0.10.0) ^61^. Data preprocessing was done in R, using limma and edgeR packages, as follows: genes were removed when weakly expressed or associated to noninformative features (“no_feature”,“ambiguous”, “too_low_aQual”,“not_aligned”, “alignment_not_unique”); and for features without as least 1 read per million in 3 of the samples (minimum number of replicates). After applying these quality criteria, we obtained 16 152 expressed genes for downstream analysis. The counts per gene were normalized to counts per million (CPMs) by dividing it by the total number of mapped reads per sample and multiplying by 10^6. The CPM normalized data were then transformed with log2 using an offset of 1. Pairwise analysis of differential gene expression was performed using the generalized linear model workflow ^62^ and the cutoffs |log2(FC)|>1 and FDR<0.05. Row z-scores for each gene represented in the heatmaps were calculated using log2 (CPMs+1) values and using heatmap.2 function from gplots package in R.

#### Alternative splicing analysis

For AS event-level analysis, raw reads were mapped to previously annotated AS events for the mouse reference assembly (mm10 vastdb.mm2.23.06.20) from VastDB (VastDB v2 released). Abundance of AS events in percent spliced in (PSI) values was estimated using VAST-TOOLS (version 2.5.1) *align* and *combine* commands ^28, 29^.

AS events in which the read coverage based on corrected reads (quality score 2) does not meet the minimum threshold (N) and those in which PSI could not be determined (NA) in at least 1 sample, and those in which minimum number of reads<10 in at least 3 samples were excluded from subsequent analysis. To calculate differential AS between each pair of time- points for the AS events, we used Vast-tools *diff* command. Vast *diff* uses Bayesian inference followed by differential analysis to calculate ΔPSI and minimum value difference (MV) between 2 samples for each AS event. We used a confidence interval of 95%, MV>=10% (**Table S1**). After these criteria, we obtained 460 events identified as being differentially AS in at least one pairwise comparison between time-points, associated with 355 genes.

To analyse the expression levels of genes undergoing differential AS in at least one pairwise comparison between time-points, we divided the 16 152 expressed genes into 4 bins of equal size according to their absolute level of expression (CPMs) (low expression, medium-low expression, medium-high expression, and high expression). Then, we distributed the genes undergoing differential AS in at least one pairwise comparison between time-points (355 genes) within these bins.

To select genes that undergo differential AS in at least one pairwise comparison between time-points but do not undergo changes in gene expression in the same time interval, we used a confidence interval of 95% with MV>=10% (differential AS) and |log2(FC)|>1 with FDR<0.05 (differential gene expression). After this cutoff we obtained 371 AS events associated with 295 genes.

Clustering of AS events into distinct sets based on their kinetics along lung development, was performed by two methods: K-means and hierarchical clustering. K-means clustering was performed using the *kmeans* function (amap package) in R using Spearman correlation as a correlation method for each pair of AS events based on their PSI values at the different time- points. The average silhouette width method was used to find the best number of clusters, using the *fviz_nbclust* function (factoextra package) in R. The AS events included in each cluster are listed in **Table S5**. Hierarchical clustering was performed using *hclust* function (stats package) in R using Spearman correlation as a correlation method for each pair of AS events based on their PSI values at the different time-points and the complete-linkage method as clustering method.

Heatmaps of clustered AS events were represented in heatmaps using the heatmap.2 function in R. For the heatmap representation, logit (PSI) values were used and were scaled by row by calculating row z-scores.

For transcript-level analysis, transcripts isoform abundance in RNAseq datasets in transcripts per million (TPM) was estimated using Kallisto (version 0.44.0) ^37^. Vegfa *i5* isoform was manually annotated in Kallisto index built with reference transcriptome before performing the pseudo-alignment.

#### Pathway analysis

KEGG pathway analysis was carried out using DAVID v6.8 ^63, 64^. KEGG terms with a modified Fisher Exact p-value (EASE score) < 0.05 and FDR<0.1 were considered. The set of 16152 genes we identified to be expressed in the lungs during the time-points analysed was used as control background list.

#### RBPs motifs enrichment analysis

To identify potential RPBs regulators of the AS events detected during lung development, we used Matt, a unix toolkit that searches for RBPs motifs in genomic sequences ^32^. Briefly, we used the *get_vast* command to extract subsets of reported AS events generated by *Vast-tools.* We considered only alternative splicing (AS) events in non-differentially expressed genes. Then we filtered the table for intron retention events (IR) and exon skipping events of type S, C1 and C2, with PSI values with a minimum quality flag of LOW. We defined categories for each event (enhanced or silenced) based on the ΔPSI values obtained from Vast-tools (confidence interval of 95% with MV>=10%). A set of unregulated events was randomly selected from the same genes with significant AS events to be used as control in the enrichment analysis. Upstream and downstream sequences of each event were obtained with Matt’s *get_seqs* command, comprising the 250 nucleotides flanking each splice site (150 nt towards the intron and 100 nt towards the exon). Then, we performed an enrichment test using Matt’s test_cisbp_enrich function, which compares the positional density of motifs from the database ^32^ in enhanced or silenced AS events against the background set of unregulated exons or introns, predicting the number of hits between the two groups of sequences associated with each rna binding protein (RBP) motif ^33^. The Matt function performs a permutation test to determine significant enrichment/depletion of RBP motifs (p- value < 0.005). Finally, we used a custom python script based on the *regex* package to find the number of events containing hits for each RBP motif, and plotted these values normalized by the total number of events of each type. In addition, we used Matt’s function get_regexp_prof to plot the positional distributions of motif hits across sequences.

We selected RBPs that undergo changes in gene expression between E18.5 and P5 (|log2FC|>1 and FDR<0.05).

To reinforce the results of the putative regulators we explored the CLIP-seq profiles of 356 RBPs in human cancer cells (HepG2 and K562), previously published and available in ENCODE ^65^. First, mouse AS events were converted to the respective human homologous exons using VastDB ^28^, and the equivalent genomic intervals (250nt flanking each splice site) were crossed with the different CLIP-seq profiles for the enriched RBPs. Individual CLIP-seq profiles for each loci were produced using Bedtools ^66^ and R tool ^67^.

### Statistical analysis

Statistical analysis was performed using GraphPad Prism 8. Measurements were taken from distinct samples, and statistical details of experiments are reported in the figures and figure legends. Sample size is reported in the figure legends. The biological replicate is defined as the number of cells, images, animals, as stated in the figure legends. Comparisons between two experimental groups were analysed with two-tailed unpaired t-test, while multiple comparisons between more than two experimental groups were assessed with one-way ANOVA with Tukey correction for multiple comparisons. Unless otherwise specified, data are represented as mean ± SD; n.s. - p > 0.05; * - p < 0.05, ** - p < 0.01, *** - p < 0.001, and **** - p < 0.0001. We considered a result significant when p<0.05.

## References

1. Mammoto, A. & Mammoto, T. Vascular Niche in Lung Alveolar Development, Homeostasis, and Regeneration. Front. Bioeng. Biotechnol. 7, 1–16 (2019).

2. Hogan, B. L. M. et al. Repair and regeneration of the respiratory system: Complexity, plasticity, and mechanisms of lung stem cell function. Cell Stem Cell 15, 123–138 (2014).

3. Vila Ellis, L. & Chen, J. A cell-centric view of lung alveologenesis. Dev. Dyn. (2020) doi:10.1002/dvdy.271.

4. Kina, Y. P., Khadim, A., Seeger, W. & El Agha, E. The Lung Vasculature: A Driver or Passenger in Lung Branching Morphogenesis? Front. Cell Dev. Biol. 8, 1–7 (2021).

5. Zepp, J. A. et al. Genomic, epigenomic, and biophysical cues controlling the emergence of the lung alveolus. Science (80-.). 371, (2021).

6. Ellis, L. V., et al. Epithelial Vegfa specifies a distinct endothelial population in the mouse lung. *bioRxiv* 52, (2019).

7. Yamamoto, H. et al. Epithelial-vascular cross talk mediated by VEGF-A and HGF signaling directs primary septae formation during distal lung morphogenesis. Dev. Biol. 308, 44–53 (2007).

8. Vila Ellis, L., et al. Epithelial Vegfa Specifies a Distinct Endothelial Population in the Mouse Lung. Dev. Cell 52, 617–630.e6 (2020).

9. Thébaud, B. et al. Vascular endothelial growth factor gene therapy increases survival, promotes lung angiogenesis, and prevents alveolar damage in hyperoxia-induced lung injury: Evidence that angiogenesis participates in alveolarization. Circulation 112, 2477–2486 (2005).

10. Farini, D. et al. A Dynamic Splicing Program Ensures Proper Synaptic Connections in the Developing Cerebellum. Cell Rep. 31, 107703 (2020).

11. Baralle, F. E. & Giudice, J. Alternative splicing as a regulator of development and tissue identity. Nat. Rev. Mol. Cell Biol. 18, 437–451 (2017).

12. Weyn-Vanhentenryck, S. M. et al. Precise temporal regulation of alternative splicing during neural development. Nat. Commun. 9, (2018).

13. Brinegar, A. E. et al. Extensive alternative splicing transitions during postnatal skeletal muscle development are required for calcium handling functions. Elife 6, (2017).

14. Treutlein, B. et al. Reconstructing lineage hierarchies of the distal lung epithelium using single-cell RNA-seq. Nature 509, 371–375 (2014).

15. Wang, Y. et al. Pulmonary alveolar type I cell population consists of two distinct subtypes that differ in cell fate. Proc. Natl. Acad. Sci. U. S. A. 115, 2407–2412 (2018).

16. Ng, Y. S., Rohan, R., Sunday, M. E., Demello, D. E. & D’Amore, P. A. Differential expression of VEGF isoforms in mouse during development and in the adult. Dev. Dyn. 220, 112–121 (2001).

17. Greenberg, J. M. et al. Mesenchymal Expression of Vascular Endothelial Growth Developing Lung. 153, 144–153 (2002).

18. Healy, A. M., Morgenthau, L., Zhu, X. & Farber, H. W. VEGF Is Deposited in the Subepithelial Matrix at the Leading Edge of Branching Airways and Stimulates Neovascularization in the Murine Embryonic Lung. 352, 341–352 (2000).

19. Domigan, C. K. et al. Autocrine VEGF maintains endothelial survival through regulation of metabolism and autophagy. J. Cell Sci. 128, 2236–2248 (2015).

20. Peach, C. J. et al. Molecular pharmacology of VEGF-A isoforms: Binding and signalling at VEGFR2. Int. J. Mol. Sci. 19, (2018).

21. Yamamoto, H., Rundqvist, H., Branco, C. & Johnson, R. S. Autocrine VEGF isoforms differentially regulate endothelial cell behavior. Front. Cell Dev. Biol. 4, 1–12 (2016).

22. Bowler, E. & Oltean, S. Alternative splicing in angiogenesis. Int. J. Mol. Sci. 20, (2019).

23. Galambos, C. et al. Defective pulmonary development in the absence of heparin- binding vascular endothelial growth factor isoforms. Am. J. Respir. Cell Mol. Biol. 27, 194–203 (2002).

24. Compernolle, V. et al. Loss of HIF-2α and inhibition of VEGF impair fetal lung maturation, whereas treatment with VEGF prevents fatal respiratory distress in premature mice. Nat. Med. 8, 702–710 (2002).

25. Beauchemin, K. J. et al. Temporal dynamics of the developing lung transcriptome in three common inbred strains of laboratory mice reveals multiple stages of postnatal alveolar development. (2016) doi:10.7717/peerj.2318.

26. LungMAP. http://www.lungmap.net.

27. Domingo-Gonzalez, R. et al. Diverse homeostatic and immunomodulatory roles of immune cells in the developing mouse lung at single cell resolution. Elife 9, 1–39 (2020).

28. Tapial, J. et al. An atlas of alternative splicing profiles and functional associations reveals new regulatory programs and genes that simultaneously express multiple major isoforms. 1759–1768 (2017) doi:10.1101/gr.220962.117.

29. Irimia, M. et al. A highly conserved program of neuronal microexons is misregulated in autistic brains. Cell 159, 1511–1523 (2014).

30. Gardina, P. J. et al. Alternative splicing and differential gene expression in colon cancer detected by a whole genome exon array. BMC Genomics 7, 325 (2006).

31. Grosso, A. R. et al. Tissue-specific splicing factor gene expression signatures. Nucleic Acids Res. 36, 4823–4832 (2008).

32. Ray, D. et al. A compendium of RNA-binding motifs for decoding gene regulation. Nature 499, 172–177 (2013).

33. Gohr, A. & Irimia, M. Matt: Unix tools for alternative splicing analysis. Bioinformatics 35, 130–132 (2019).

34. Yu, M. et al. Lack of Bcr and Abr Promotes Hypoxia-Induced Pulmonary Hypertension in Mice. PLoS One 7, e49756 (2012).

35. Tian, Y. et al. Quantitative proteomic characterization of lung tissue in idiopathic pulmonary fibrosis. Clin. Proteomics 16, 6 (2019).

36. Eichmann, A. & Simons, M. VEGF signaling inside vascular endothelial cells and beyond. Curr. Opin. Cell Biol. 24, 188–193 (2012).

37. Bray, N. L., Pimentel, H., Melsted, P. & Pachter, L. Near-optimal probabilistic RNA-seq quantification. Nat. Biotechnol. 34, 525–527 (2016).

38. Gillich, A. et al. Capillary cell-type specialization in the alveolus. Nature 586, 785–789 (2020).

39. Yang, J. et al. The development and plasticity of alveolar type 1 cells. Dev. 143, 54–65 (2016).

40. Koh, C. M. Chapter Sixteen - Preparation of Cells for Microscopy using Cytospin. In Laboratory Methods in Enzymology: Cell, Lipid and Carbohydrate (ed. Lorsch, J. B. T.-M. in E.) vol. 533 235–240 (Academic Press, 2013).

41. Hasegawa, K. et al. Fraction of MHCII and EpCAM expression characterizes distal lung epithelial cells for alveolar type 2 cell isolation. Respir. Res. 18, 1–13 (2017).

42. Raredon, M. S. B. et al. Single-cell connectomic analysis of adult mammalian lungs. Sci. Adv. 5, 1–16 (2019).

43. Coomer, A. O., Black, F., Greystoke, A., Munkley, J. & Elliott, D. J. Alternative splicing in lung cancer. Biochim. Biophys. acta. Gene Regul. Mech. 1862, 194388 (2019).

44. Kusko, R. L. et al. Integrated Genomics Reveals Convergent Transcriptomic Networks Underlying Chronic Obstructive Pulmonary Disease and Idiopathic Pulmonary Fibrosis. Am. J. Respir. Crit. Care Med. 194, 948–960 (2016).

45. Calderone, V. et al. Sequential Functions of CPEB1 and CPEB4 Regulate Pathologic Expression of Vascular Endothelial Growth Factor and Angiogenesis in Chronic Liver Disease. Gastroenterology 150, 982–997.e30 (2016).

46. Pascale, A., Amadio, M. & Quattrone, A. Defining a neuron: Neuronal ELAV proteins. Cell. Mol. Life Sci. 65, 128–140 (2008).

47. Kato, Y. et al. ELAVL2-directed RNA regulatory network drives the formation of quiescent primordial follicles. EMBO Rep. 20, e48251 (2019).

48. Liu, T.-Y. et al. Muscle developmental defects in heterogeneous nuclear Ribonucleoprotein A1 knockout mice. Open Biol. 7, (2017).

49. Kato, K. et al. Pulmonary pericytes regulate lung morphogenesis. Nat. Commun. 9, 1– 14 (2018).

50. Nantie, L. B. et al. Lats1/2 inactivation reveals hippo function in alveolar type i cell differentiation during lung transition to air breathing. Dev. 145, (2018).

51. Lee, S. et al. Autocrine VEGF Signaling Is Required for Vascular Homeostasis. 691–703 (2007) doi:10.1016/j.cell.2007.06.054.

52. Krock, B. L., Skuli, N. & Simon, M. C. Hypoxia-induced angiogenesis: good and evil. Genes Cancer 2, 1117–1133 (2011).

53. Yue, L., Wan, R., Luan, S., Zeng, W. & Cheung, T. H. Dek Modulates Global Intron Retention during Muscle Stem Cells Quiescence Exit. Dev. Cell 53, 661–676.e6 (2020).

54. Jacob, A. G. & Smith, C. W. J. Intron retention as a component of regulated gene expression programs. Hum. Genet. 136, 1043–1057 (2017).

55. Wong, J. J. L., Au, A. Y. M., Ritchie, W. & Rasko, J. E. J. Intron retention in mRNA: No longer nonsense: Known and putative roles of intron retention in normal and disease biology. BioEssays 38, 41–49 (2016).

56. Mauger, O., Lemoine, F. & Scheiffele, P. Targeted Intron Retention and Excision for Rapid Gene Regulation in Response to Neuronal Activity. Neuron 92, 1266–1278 (2016).

57. Bentley, D. L. Coupling mRNA processing with transcription in time and space. Nat. Rev. Genet. 15, 163–175 (2014).

58. Pfaffl, M. W. A new mathematical model for relative quantification in real-time RT- PCR. Nucleic Acids Res. 29, e45 (2001).

59. Kim, D. et al. TopHat2: accurate alignment of transcriptomes in the presence of insertions, deletions and gene fusions. Genome Biol. 14, R36 (2013).

60. Langmead, B. & Salzberg, S. L. Fast gapped-read alignment with Bowtie 2. Nat. Methods 9, 357–359 (2012).

61. Anders, S., Pyl, P. T. & Huber, W. HTSeq--a Python framework to work with high- throughput sequencing data. Bioinformatics 31, 166–169 (2015).

62. Robinson, M. D., McCarthy, D. J. & Smyth, G. K. edgeR: a Bioconductor package for differential expression analysis of digital gene expression data. Bioinformatics 26, 139–140 (2010).

63. Huang, D. W., Sherman, B. T. & Lempicki, R. A. Bioinformatics enrichment tools: paths toward the comprehensive functional analysis of large gene lists. Nucleic Acids Res. 37, 1–13 (2009).

64. Huang, D. W., Sherman, B. T. & Lempicki, R. A. Systematic and integrative analysis of large gene lists using DAVID bioinformatics resources. Nat. Protoc. 4, 44–57 (2009).

65. “A large-scale binding and functional map of human RNA-binding proteins | Nature.” https://www.nature.com/articles/s41586-020-2077-3.

66. Quinlan, A. R. & Hall, I. M. BEDTools: A flexible suite of utilities for comparing genomic features. Bioinformatics 26, 841–842 (2010).

67. https://www.r-project.org/. “R: The R Project for Statistical Computing.”.https://www.r-project.org/.

